# Diversity and assembly of the microbiome of a leguminous plant along an urbanization gradient

**DOI:** 10.1101/2024.06.10.598260

**Authors:** David Murray-Stoker, Laura Rossi, Marc T. J. Johnson

**Affiliations:** Department of Ecology and Evolutionary Biology, University of Toronto, Toronto, Ontario, Canada; Department of Biology, University of Toronto Mississauga, Mississauga, Ontario, Canada; Centre for Urban Environments, University of Toronto Mississauga, Mississauga, Ontario, Canada; Farncombe Family Digestive Health Research Institute, McMaster University, Hamilton, Ontario, Canada; Department of Medicine, McMaster University, Hamilton, Ontario, Canada

**Keywords:** Biodiversity, community assembly, legume-rhizobia, plant microbiome, soil microbiome, urban ecology

## Abstract

Interactions between plants and bacterial communities are essential for host physiology and broader ecosystem functioning, but plant-microbiome interactions can be disrupted by environmental change like urbanization. Here, we evaluated how urbanization affected the diversity and assembly of soil and white clover (*Trifolium repens*) microbiome communities. We sampled 35 populations of white clover and associated roots and soil along an urbanization gradient. Soil alpha diversity was greater at the urban and rural limits of the gradient and lower in suburban habitats, while root alpha diversity was not influenced by urbanization. Root and soil bacterial communities had distinct compositions, with greater beta diversity for root compared to soil microbiomes. We found that urbanization directly and indirectly affected soil microbiome assembly, particularly through soil carbon. In contrast, root microbiome assembly was not linked to urbanization, which suggested that the host plant acted as an additional filter on microbiome assembly independent of urbanization. We also found that key pathogenic bacteria like *Legionella* and *Clostridium* varied in abundance with urbanization, which has implications for human health. Together, our study underscores the importance of examining how urban-driven environmental change alters the ecology and function of soil and root microbiomes.

## Introduction

Plants are associated with a vast diversity of bacteria that are essential for the health and physiology of their hosts [1–3]. Bacteria comprising the plant microbiome are also important for ecological and evolutionary processes, such as nutrient acquisition and cycling [4], responses to abiotic stressors [e.g., drought, climate change; 5,6], and adaptation to novel environments [7,8]. Plants assemble their root microbiomes from the surrounding soil, where the soil acts as a pool from which bacteria are filtered and recruited by the host plant [2–4,9]. Alterations to the soil microbiome can therefore have cascading effects on plant root microbiome assembly and function. Given the ecological and evolutionary importance of plant microbiomes, it is essential to understand how ongoing environmental change affects microbiome assembly and diversity [10,11].

Urbanization is among the most dominant forms of anthropogenic disturbance and environmental change on Earth [12]. The process of urbanization frequently alters the biotic and abiotic environment, including increased impervious surface cover, elevated pollution (e.g., light, noise, nutrient), and altered community composition and diversity[13–16]. Spatial variation in soil chemistry and nutrients driven by urbanization can affect the composition of soil microbial communities [17–19], which selects for bacteria that can tolerate the environmental conditions and, in turn, determines the assembly of plant microbiomes [2–4,9,20,21]. Previous research has found positive [22,23], negative [24], and neutral [25] effects of urbanization on soil bacterial community diversity. Soil bacterial community composition can vary depending on the habitat type, such as between large parks and road medians [18] or between suburban and urban forests [23]. There is also emerging evidence that urbanization can affect the distribution of mutualistic and pathogenic bacteria [24], which can have implications for both plant and human health [26].

Despite considerable research on the diversity and composition of soil bacterial communities in urban environments [17–19,22–25,27], there is little research evaluating how plant microbiome assembly and diversity is affected by urban-driven environmental change [but see 28–30].

Here, we evaluated plant microbiome assembly and diversity of white clover (*Trifolium repens*) along an urbanization gradient. A previous study found that urbanization altered the dynamics of the mutualism between white clover and rhizobia [31], whereby urban plants had reduced investment in rhizobia and urbanization affected the source of nitrogen used by white clover. It was also found that soil nitrogen was the key link between urbanization and the mutualistic interaction [31]; however, the relative importance of the broader bacterial microbiome on the ecology of the mutualism was not evaluated. In the present study, we asked three specific questions: (Q1) How does the diversity and composition of the microbiome vary with urbanization? (Q2) What are the direct and indirect effects of urbanization on soil chemistry and microbiome assembly? And (Q3) how does the prevalence of mutualistic, pathogenic, and ecosystem function bacteria change with urbanization? By quantifying the changes to microbiome structure along an urbanization gradient, we can examine the community- and ecosystem-level consequences of urbanization on plant microbiomes more comprehensively.

## Methods

### Study System

We used white clover (*Trifolium repens* L., Fabaceae) as a model host organism to study plant-microbiome interactions along an urbanization gradient [16]. White clover is a perennial, herbaceous legume that reproduces clonally through stolons and sexually via obligately-outcrossed flowers [32]. Native to Eurasia, white clover has expanded globally following its introduction as livestock fodder and as a cover crop [32,33], including the study area of the Greater Toronto Area, Canada. White clover interacts in a facultative mutualism with *Rhizobium leguminosarum* symbiovar *trifolii* (hereafter rhizobia), its primary rhizobial symbiont [34]. Rhizobia typically comprise the majority of the bacterial microbiome in legume roots [20,21,35].

### Field Sampling

White clover, associated roots, and soil samples were collected at 35 sites along an urbanization gradient in July 2020 (figure 1). At each site (hereafter population), we collected 5 white clover individuals and their associated roots and nodules, with plants separated by a minimum of 5 m to avoid collecting the same clone. Five soil cores were also collected at each population, with each core taken ∼5 m from a sampled white clover individual; we also ensured soil cores avoided conspecific plants and other nitrogen-fixing legumes (e.g., *Medicago lupulina*). Soil cores were taken to a depth of 5 cm and we removed the top 2 cm of organic material because it could confound soil nutrient analyses. Individual soil cores were pooled for a composite soil sample for each population. We stored white clover samples at −20°C and soil samples at −80°C until later processing.

**Figure 1:**
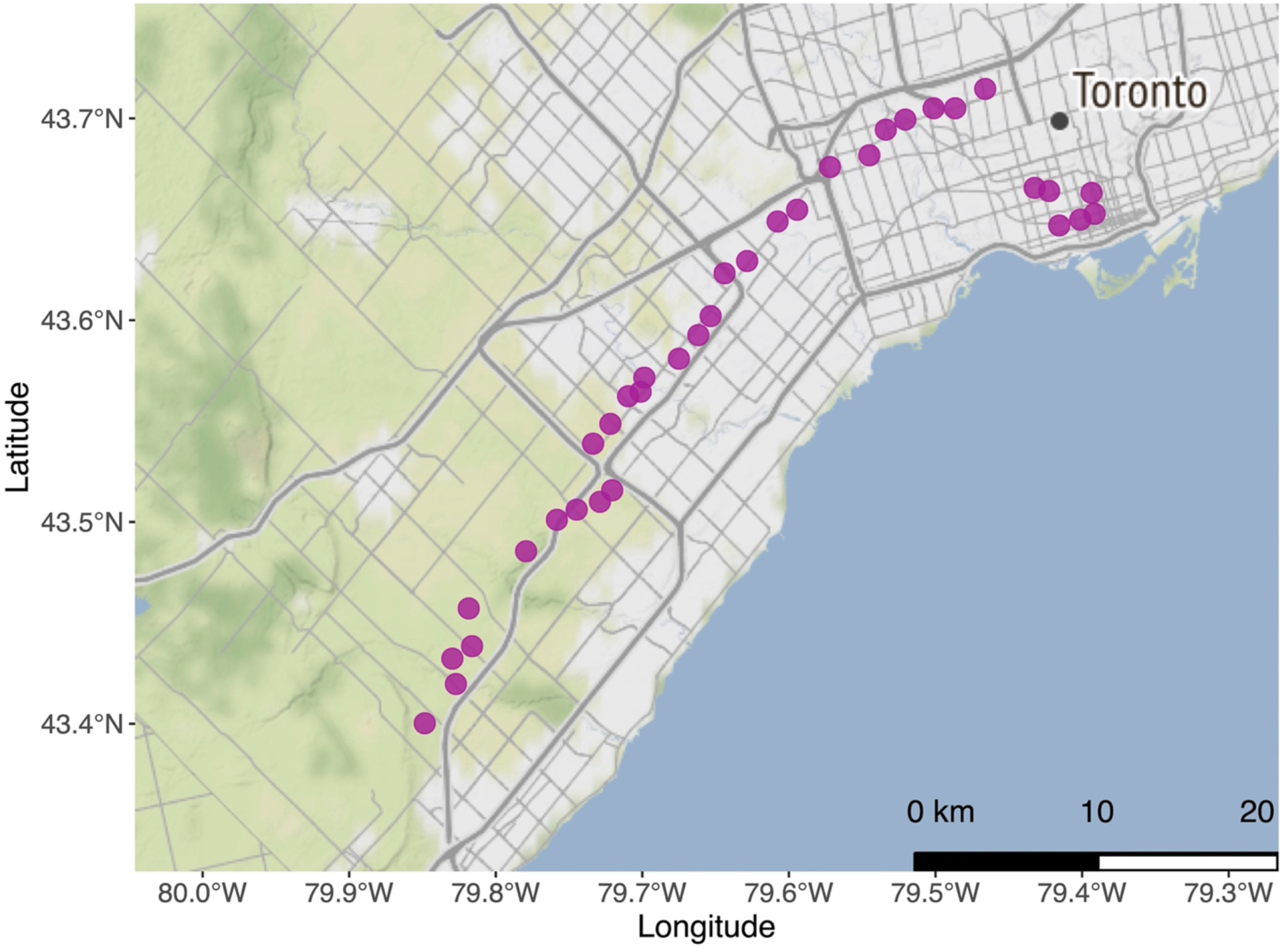
Map of the urbanization gradient in the Greater Toronto Area, Ontario, Canada with the location of the 35 focal populations indicated by purple circles. Urban and suburban areas are in gray, nonurban agricultural and forested areas in green, and Lake Ontario in light blue. Map tiles by Stamen Design, under CC BY 3.0. Data by OpenStreetMap, under ODbL.

### Carbon and Nitrogen Analyses

*Leaf processing:* We excised one fully-expanded trifoliate leaf from each white clover individual, with leaves pooled within each population to generate a composite leaf tissue sample. Leaf samples were then freeze-dried, ground, and homogenized using a tissue grinder (FastPrep 96, MP Biomedicals, Irvine, CA, USA).

*Soil processing:* Soil samples were removed from the freezer and thawed overnight in the lab (20°C). We then filtered samples over stacked sieves (mesh diameters: 4.75 mm, 2 mm, 1 mm, and 0.5 mm) to remove rocks, gravel, and plant material. Soil retained on the 0.5 mm sieve was collected and then dried at 60°C for 48 hrs. Soil samples were then freeze-dried, ground, and homogenized using a tissue grinder (FastPrep 96, MP Biomedicals, Irvine, CA, USA).

*Elemental and isotope analyses:* To evaluate the amount (%) and type (isotope natural abundances) of carbon (C) and nitrogen (N) in the soil and white clover leaves, we conducted elemental and stable isotope analyses. Leaf tissue (≈ 2 mg) and soil samples (≈ 30 mg) were weighed on a microbalance (XP2U Mettler Toledo, Mississauga, ON, Canada) and packed into tin capsules (Costech Analytical Technologies Inc., Valencia, CA, USA). All samples were analyzed for percent carbon (C) and nitrogen (N) and their respective stable isotopes (i.e., ^13/12^C, ^15/14^N) using a Carlo Erba NA 1500C/H/N Analyzer coupled to a Thermo Delta V IRMS system (Thermo Fisher Scientific, Waltham, MA, USA). Stable isotopes were calculated relative to a standard in delta (δ) notation [36]:

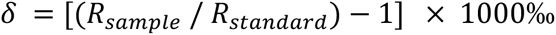

where R is the ratio of heavier-to-lighter isotopes (i.e., ^13^C to ^12^C = δ^13^C, ^15^N to ^14^N = δ^15^N). All samples were analyzed at the Stable Isotope Ecology Laboratory (University of Georgia, Georgia, USA; http://siel.uga.edu/). Increased amounts of C and N can indicate nutrient availability, white stable isotopes indicate the type and source of C and N. Soil with enriched (i.e. higher) δ^13^C typically has increased organic matter content, while white clover leaf tissue with depleted (i.e., lower) δ^13^C suggests increased photosynthetic activity [36,37]. Soil with enriched δ^15^N can indicate anthropogenic disturbance from N deposition and septic waste [36,37]. White clover leaf tissue with depleted δ^15^N suggests increased N-fixation from rhizobia, while enriched δ^15^N indicates N acquisition directly from the soil [36–38].

### Landscape Metrics

We calculated three measures of urbanization for each population. Distance from the city centre was calculated as the distance from the population to Toronto City Hall (43.651536°N, −79.383276°W), with the distance on the ellipsoid calculated using the ‘geospher’ package [39]. We calculated the Human Influence Index (HII) for each population using a 1 km resolution raster dataset [40]. Finally, we estimated the percent impervious surface cover (ISC) within a 500 m buffer of each population using the 30 m resolution Global Man-Made Impervious Surface raster dataset [41]. Urbanization metric calculations were measured using the ‘sf’ [42] and ‘terr’ [43] packages.

### Sample Preparation and DNA Extraction

We collected all root material for each sampled white clover individual. Samples were placed in 15 mL Falcon tubes and then washed in autoclaved MilliQ water while vortexing at maximum speed for 10 s; samples were washed 3 times, with the water replaced between washes. We also sonicated root samples for 10 min at 60 Hz to facilitate physical removal of any remaining soil. After washing and sonication, roots were freeze-dried (Freeze Dryer Epsilon 2-6D LSCplus) and then ground to a fine powder (QIAGEN TissueLyser II). We weighed approximately 40 mg of roots for each root sample (mean ± SE = 43.340 mg ± 1.951 mg), with samples stored in the freezer at −80°C until DNA extraction.

Immediately prior to DNA extraction, soil samples were removed from a −80°C freezer and thawed for ∼12 hours in a refrigerator at 4°C. Samples were then freeze-dried (Freeze Dryer Epsilon 2-6D LSCplus) and then homogenized (QIAGEN TissueLyser II). Approximately 250 mg (wet weight) of each soil sample was weighed for DNA extraction (246.400 mg ± 0.049 mg). We extracted DNA from all root and soil samples using the QIAGEN DNeasy PowerSoil Pro Kit following the manufacturer’s protocol.

### Sequencing

We sought to comprehensively sequence the bacterial microbiome. To accomplish this, we targeted the hypervariable V4 region of the 16S rRNA gene using dual-indexed primers [primers 515f and 806r; 43,44], with peptide nucleic acid (PNA) clamps to inhibit amplification of plant plastid and mitochondrial DNA [46,47]. To prepare samples for sequencing, we quantified extracted DNA using a Nanodrop spectrophotometer. Soil DNA was normalized to 50 ng/µL (50.027 ng/µL ± 2.987 ng/µL), but root DNA was not normalized due to low and variable concentrations of extracted DNA (28.072 ng/µL ± 13.959 ng/µL) that could have been confounded by both microbial and plant DNA. All samples were amplified and sequenced by the Surette Lab at McMaster University on an Illumina MiSeq with 2 x 300 paired-end reads, with separate sequencing runs for soil and root samples.

### Bioinformatic Pipeline

Demultiplexed reads were processed using the DADA2 pipeline [package version 1.24.0; 47], and all code will be made available on Zenodo. We first examined quality profiles of the forward and reverse reads for 20 samples. We trimmed sequences for both root (forward = 240 bp, reverse = 230 bp) and soil (forward = 275 bp, reverse = 250 bp) reads when the Phred quality (Q) score dropped (i.e., Q < 30 or accuracy dropping below 99.9%). We removed the first 21 bp of forward and 20 bp of reverse reads due to low Q-scores and to remove primers. We also filtered any sequences with ambiguous nucleotide assignments, any singular instance of a Q-score < 2, and/or sequences with more than 2 expected errors [48]. Once we filtered and trimmed sequences, we used the DADA2 algorithm to estimate the error rates and then inferred bacterial taxa using DADA2 sample inference, which infers taxa as amplicon sequence variants (ASVs). Once ASVs were identified, we merged the forward and reverse reads and removed chimeras.

We ran separate DADA2 pipelines for the root and soil samples because reads were from independent sequencing runs and therefore had different error rates to be inferred by DADA2 [48].

Taxonomy was assigned to individual ASVs for both the root and soil samples using the RDP naïve Bayesian classifier [49] and the Ribosomal Database Project training set 18 [50] implemented in DADA2. We constructed a phylogenetic tree with both root and soil ASVs by performing a multiple sequence alignment using Clustal-Omega [51], calculating the distance matrix between sequences, and generating a neighbour-joining tree using the BIONJ algorithm [52,53].

Samples were further processed for analyses using the ‘phyloseq’ [54] and ‘tidyamplicons’ [55] packages. First, we only included ASVs assigned to the kingdom Eubacteria. Second, we rarefied samples to equal sequencing depth for each compartment (root = 15,000 reads, soil = 20,000 reads). Third, we removed potential contaminant ASVs from the root (126 potential contaminants from 16,985 total ASVs) and soil (19 potential contaminants from 20,566 total ASVs) samples that were identified using the ‘decontam’ package [56,57]. Finally, we merged the processed root and soil samples with the full neighbour-joining tree for analyses. Our final dataset included 27,006 ASVs from 161 root samples with (mean ± SE) 53,745 ± 1,699 reads per sample and 171 soil samples with 66,695 ± 1,898 reads per sample.

### Statistical Analyses

We provide detailed methods of our statistical analyses in the electronic supplementary material, Statistical Analyses. Briefly, Q1 was addressed by comparing alpha diversity [ASV richness, inverse Simpson (Simpson’s D^−1^), ASV evenness (inverse Simpson/richness), and Faith’s phylogenetic diversity (PD)] by measures of urbanization (distance from the city centre, HII, and mean ISC) using generalized additive mixed models [GAMMs; 57,58]. To further address Q1, we evaluated beta diversity as the differences in bacterial composition among communities by: (1) comparing compositions using principal coordinates analysis (PCoA) and PERMANOVA [60,61]; and (2) comparing the beta dispersion of community composition with urbanization [61,62]. For Q2, we used piecewise structural equation modeling (pSEM) to determine the hierarchical pathways of microbiome assembly [63–65]. Finally, for Q3 we used the Analysis of Compositions of Microbiomes with Bias Correction (ANCOMBC) to detect differentially-abundant taxa in relation to urbanization [66]. We also evaluated if the relative abundances of focal bacterial taxa across functional groups varied with urbanization, with focal taxa consisting of putative mutualistic, pathogenic, and ecosystem functioning bacteria (electronic supplementary material, Statistical Analyses and table S1). All of the above analyses were performed using R [version 4.2.3; 66] in the RStudio environment [version 2022.07.2; 67]. All packages used for data management and analysis are noted in electronic supplementary material, Statistical Analyses.

## Results

### How does the diversity and composition of the microbiome vary with urbanization?

Alpha diversity primarily varied by compartment (i.e., root vs. soil), with greater alpha diversity in the soil compared to the root compartment (electronic supplementary material, table S2, figure 2). Root alpha diversity did not vary with distance from the city centre, regardless of the alpha diversity metric (electronic supplementary material, table S2, figure 2). Soil ASV richness (P < 0.001, η^2^_P_ = 0.026, deviance = 68.8%) and Faith’s PD (P < 0.001, η^2^_P_ = 0.053, deviance = 74.3%) had peaks at the urban and rural limits of the urbanization gradient, and both declined in suburban areas between ∼5-10 km from the city centre (electronic supplementary material, table S2, figure 2). In contrast, soil Simpson’s D^−1^ peaked in rural environments at ∼35 km from the city centre (P < 0.001, η^2^_P_ = 0.129, deviance = 83.8%). We also found extensive variation along the urbanization gradient among populations for Simpson’s D^−1^ (P < 0.001, η^2^_P_ = 0.247) and evenness (P < 0.001, η^2^_P_ = 0.186). Trends were similar when urbanization was considered as HII or mean ISC (electronic supplementary material, table S2, figure S1).

**Figure 2:**
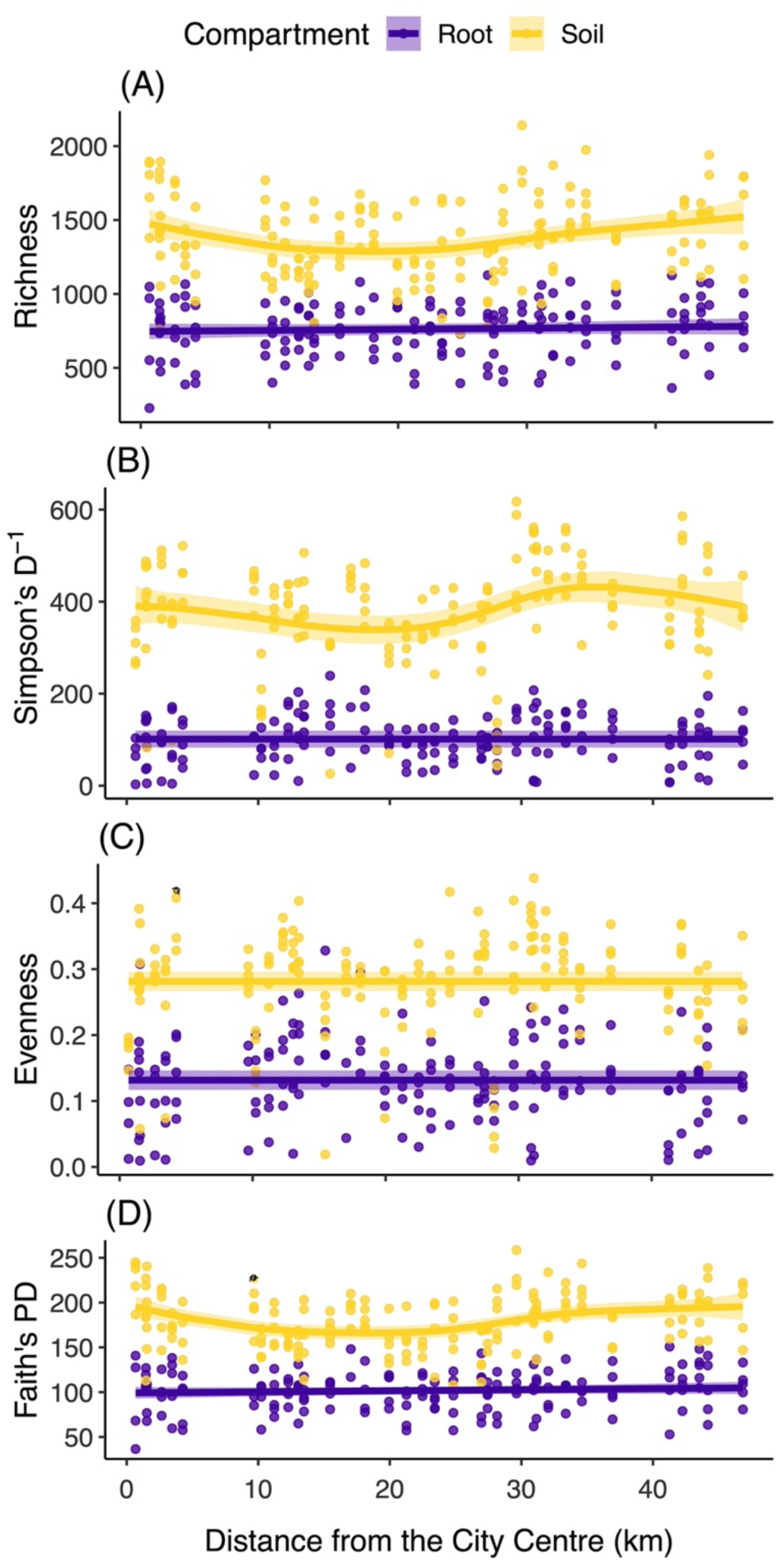
Relationship between measures of alpha diversity and distance from the city centre. Alpha diversity was quantified as richness (A), inverse Simpson (B), evenness (C), and Faith’s phylogenetic diversity (D). Points represent individual samples and lines are smoothed curves (± 95% confidence interval) from generalized additive mixed models, with separate curves shown for the root and soil microbiome compartments (purple and yellow, respectively). Deviance explained by the models: richness (68.8%), inverse Simpson (83.8%), evenness (66.1%), and Faith’s phylogenetic diversity (74.3%). Detailed statistics are provided in Table S1.

Root and soil communities had distinct bacterial composition based on the weighted UniFrac distance (figure 3), with compartment explaining 48.9% of the variation (PERMANOVA F_1,331_ = 444.480, P = 0.001). There were also effects on community composition by population (PERMANOVA F_1, 331_ = 3.616, P = 0.001, R^2^ = 0.135) and a compartment-by-population interaction (PERMANOVA F_1, 331_ = 171.431, P = 0.001, R^2^ = 0.088; figure 3). Root communities also had greater beta dispersion than soil communities (P = 0.012, η^2^_P_ = 0.033), with beta dispersion (i.e., measure of variation in bacterial composition among populations, based on weighted UniFrac dissimilarity) peaking at the urban and rural limits of the gradient and declining between ∼10-35 km (deviance = 67.2%; electronic supplementary material, table S3, figure 3). Both PERMANOVA and beta dispersion results were similar regardless of the dissimilarity matrix used to characterize community composition (i.e., Bray-Curtis, unweighted UniFrac, or weighted UniFrac). However, beta dispersion for root or soil communities only varied by distance from the city centre and not HII or mean ISC (electronic supplementary material, table S3, figures S2-S4).

**Figure 3:**
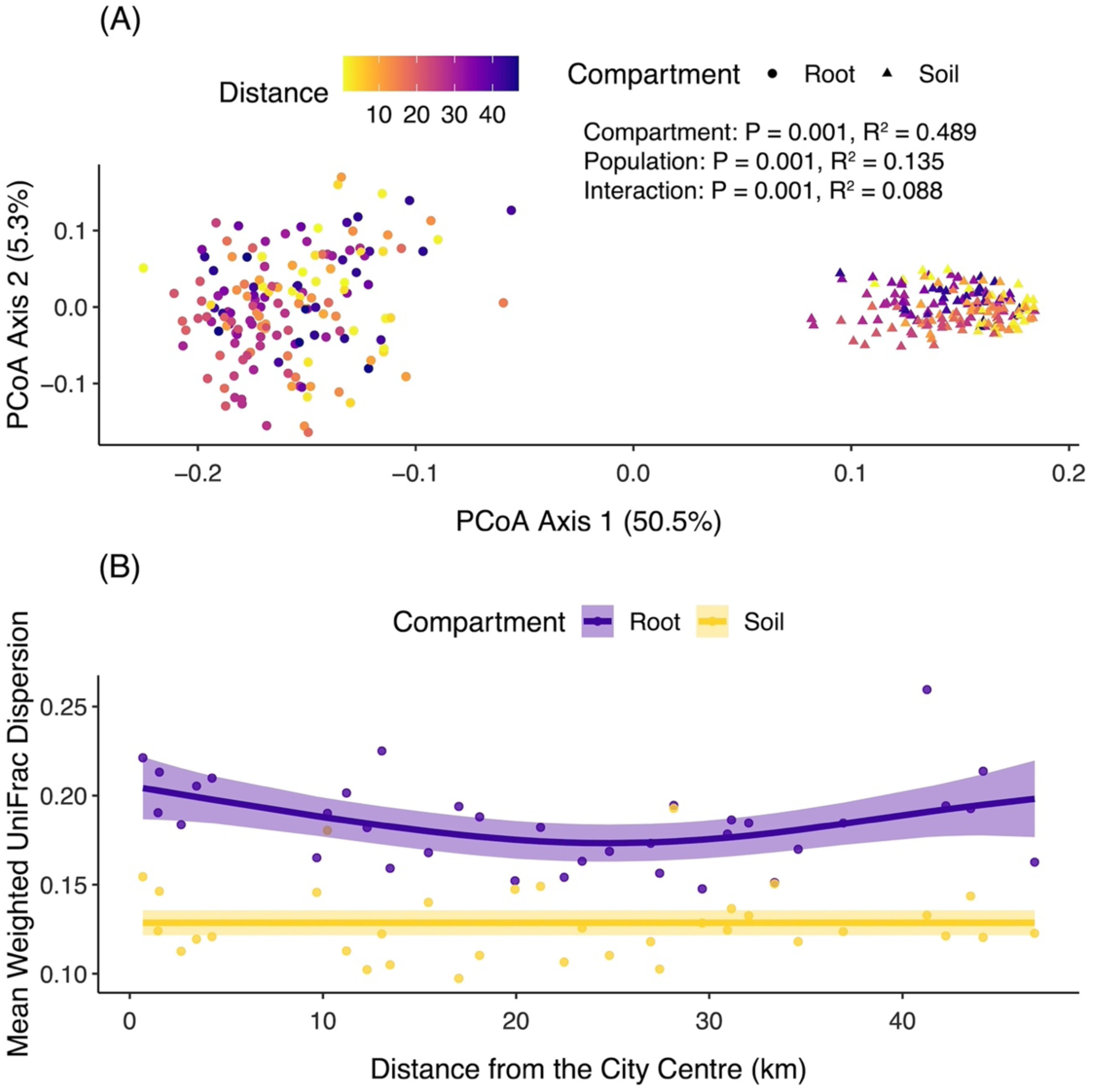
Principal coordinates analysis of the weighted UniFrac dissimilarity among bacterial communities (A), which shows distinct groupings by microbiome compartments (purple circles = root, yellow triangles = soil). Points are coloured by distance from the city centre. Inset text presents results from a PERMANOVA comparing composition by microbiome compartment, population, and their interaction. We also show the relationship between mean weighted UniFrac beta dispersion and distance from the city centre (B). Points represent population means and lines are smoothed curves (± 95% confidence interval) from a generalized additive model, with separate curves shown for the root and soil microbiome compartments (purple and yellow, respectively). Deviance explained by the model = 67.2%, with detailed statistics provided in Table S3.

### What are the direct and indirect effects of urbanization on soil chemistry and microbiome assembly?

Urbanization had indirect effects on microbiome assembly and direct effects on potential ecosystem functions (electronic supplementary material, table S4, figure 4). Soil microbiome assembly was mediated through soil % C, which was directly (path coefficient = 0.813) and indirectly influenced by HII (compound path coefficient = |−0.626| ⨉ 0.813 = 0.509); indirect effects of HII were embedded in correlational pathways between soil % C, % N, δ^13^C, and δ^15^N (electronic supplementary material, table S4, figure 4). Root microbiome assembly was influenced by soil microbiome composition but not urbanization, although root microbiome assembly further affected leaf C and N (electronic supplementary material, table S4, figure 4). Human Influence Index had a direct effect on leaf % C (path coefficient = −0.487) in addition to indirect effects mediated through soil δ^15^N on leaf % C (compound path coefficient = 0.386 ⨉ 0.426 = 0.164) and leaf % N (compound path coefficient = 0.386 ⨉ 0.544 = 0.210), while mean ISC had a direct effect on both leaf % C (path coefficient = 0.413) and leaf % N (path coefficient = 0.361).

**Figure 4:**
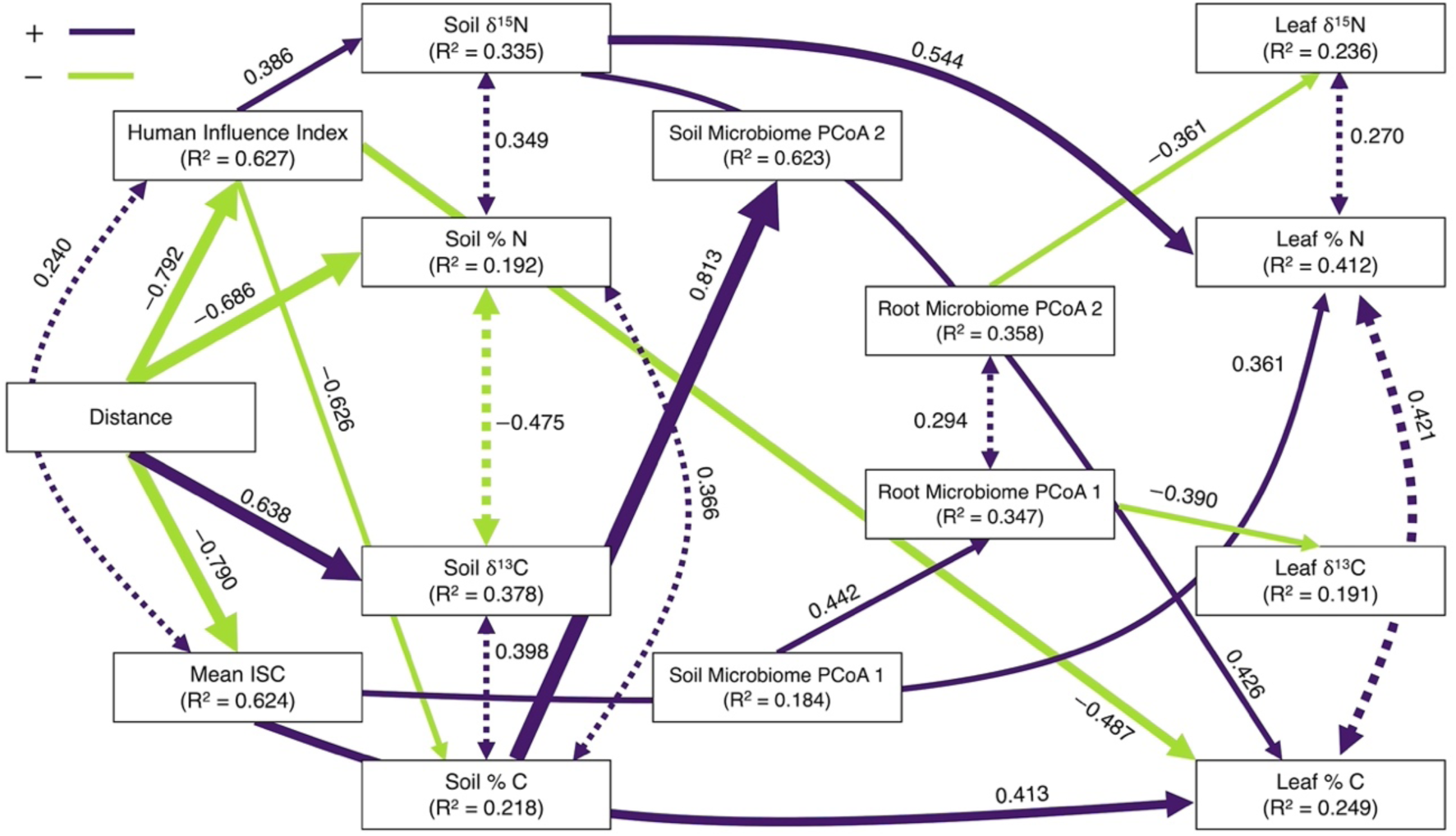
Path diagram of the hypothesized causal pathways underlying microbiome assembly (Fisher’s C = 77.027, df = 68, P = 0.212). Lines represent pathways in the model, with only pathways with P < 0.100 presented for graphical clarity. Solid single-headed arrows indicate a causal pathway and dashed double-headed arrows indicate a correlational pathway between variables. Purple lines indicate positive pathways, while green lines indicate negative pathways. Line widths are scaled relative to the magnitude of the path coefficients. The R^2^ is reported for each endogenous variable. Detailed statistics for all structural equations are provided in Table S4.

### How does the prevalence of mutualistic, pathogenic, and ecosystem function bacteria change with urbanization?

We found that ∼17% of all bacteria (71/409 genera) had differential abundance in relation to distance from the city center, although abundances more strongly differed between compartments than with urbanization (figure 5). Abundances were typically greater in the soil compared to the root compartment (mean log-fold change (LFC) ± SE = −0.156 ± 0.218), notwithstanding considerable variation in both direction and magnitude of the LFC among genera (figure 5). While abundances typically increased with increasing distance from the city centre (LFC = 0.003 ± 0.002; figure 5), there was also a weak (LFC = −0.0001 ± 0.010) and infrequent (∼2% of genera) interaction between compartment and distance from the city centre, indicating that the root and soil microbiome tended to respond to urbanization in the same way (figure 5).

**Figure 5:**
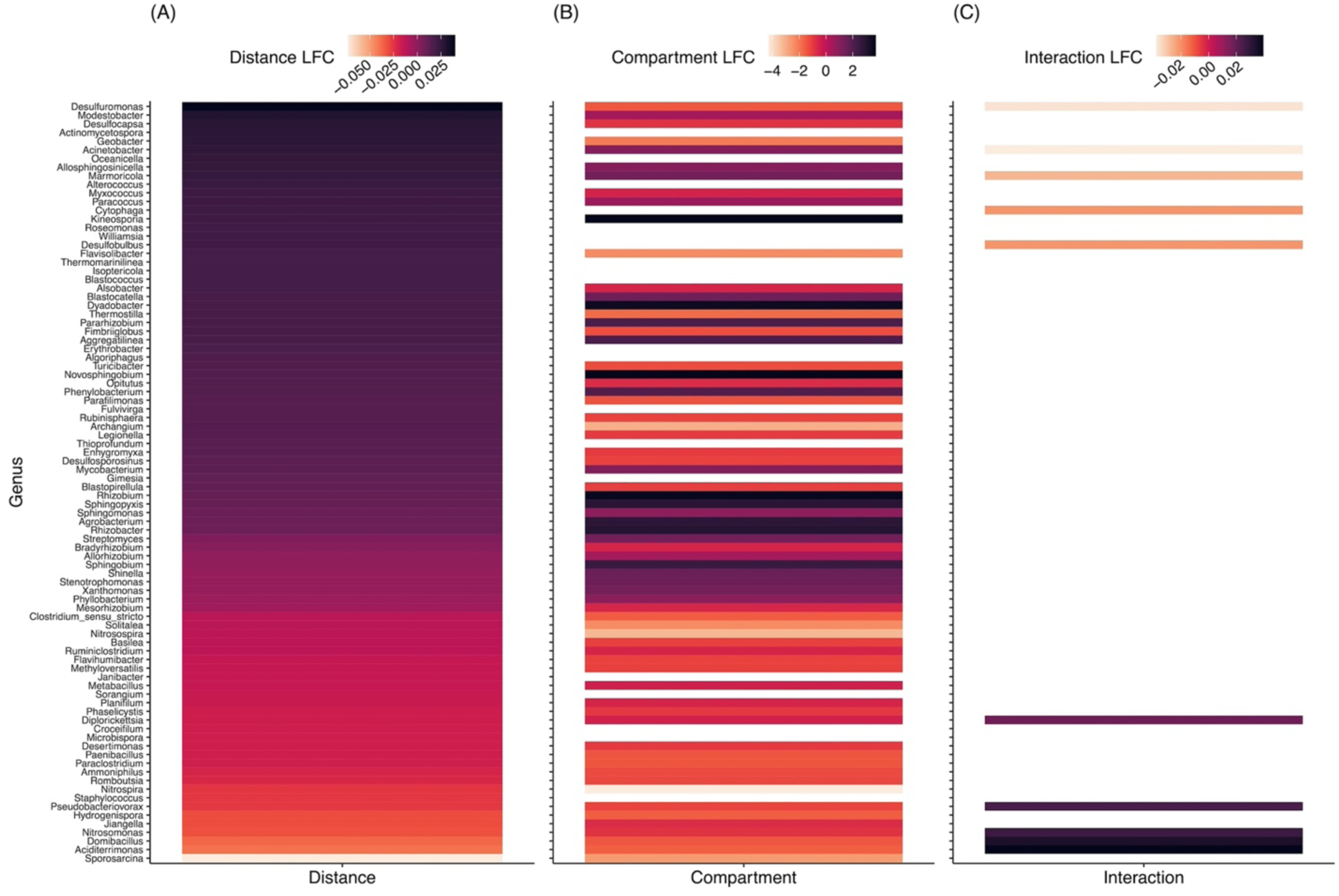
Heatmaps of differentially-abundant bacterial taxa by distance from the city centre (A), compartment (root or soil; B) and the interaction (C) following the ANCOMBC procedure. For graphical clarity, we only show genera with P < 0.100 for the effect of distance from the city center (71/409 total genera), except for any focal bacteria that had P > 0.100 (17/33 focal genera). Taxa are presented in descending log-fold change (LFC) in relation to distance from the city centre. Positive LFCs indicate increased abundance with increasing distance (A), increased abundance in the root compartment (B), and increased abundance in the root compartment with increasing distance (C). Full results of the differential abundance analysis are provided in the electronic supplementary material. Note: the LFC scale is different for each panel.

Relative abundances of mutualistic, pathogenic, and ecosystem functioning bacteria varied by compartment and urbanization, but effects depended on bacteria identity (figure 5). Mutualistic bacteria primarily varied by compartment (electronic supplementary material, table S5, figure 6), although *Bradyrhizobium* and *Rhizobium* also varied with urbanization in the root (*Rhizobium*) or soil (*Bradyrhizobium*) compartments (electronic supplementary material, table S5, figure 6). *Bradyrhizobium* in the soil had lower relative abundance at the urban and rural limits of the urbanization gradient, peaking in relative abundance in suburban areas ∼25 km from the city centre and just before the transition to rural areas (electronic supplementary material, table S5, figure 6). In contrast, *Rhizobium* relative abundance rapidly declined in the root from its peak in the urban limit, with lowest abundances in the suburbs 10-20 km from the city centre and abundances then progressively increased with increasing distance from the city centre (electronic supplementary material, table S5, figure 6). *Agrobacterium*, a plant pathogen, increased in relative abundance in the root with increasing distance from the city centre, particularly at the rural limit (electronic supplementary material, table S6, figure 6). In contrast, *Nitrososomas* and *Nitrospira* − bacteria involved in the nitrogen cycle − had greater relative abundances in the soil in the urban limit of the gradient (electronic supplementary material, table S7, figure 6). Both ANCOMBC and focal bacteria analyses produced similar results if urbanization was quantified as HII or mean ISC (electronic supplementary material, tables S5-S7, figures S7-S9). Complete ANCOMBC and focal bacteria results are provided in the electronic supplementary material (electronic supplementary material, tables S5-S7, figures S7-S9, ANCOMBC Dataset).

**Figure 6:**
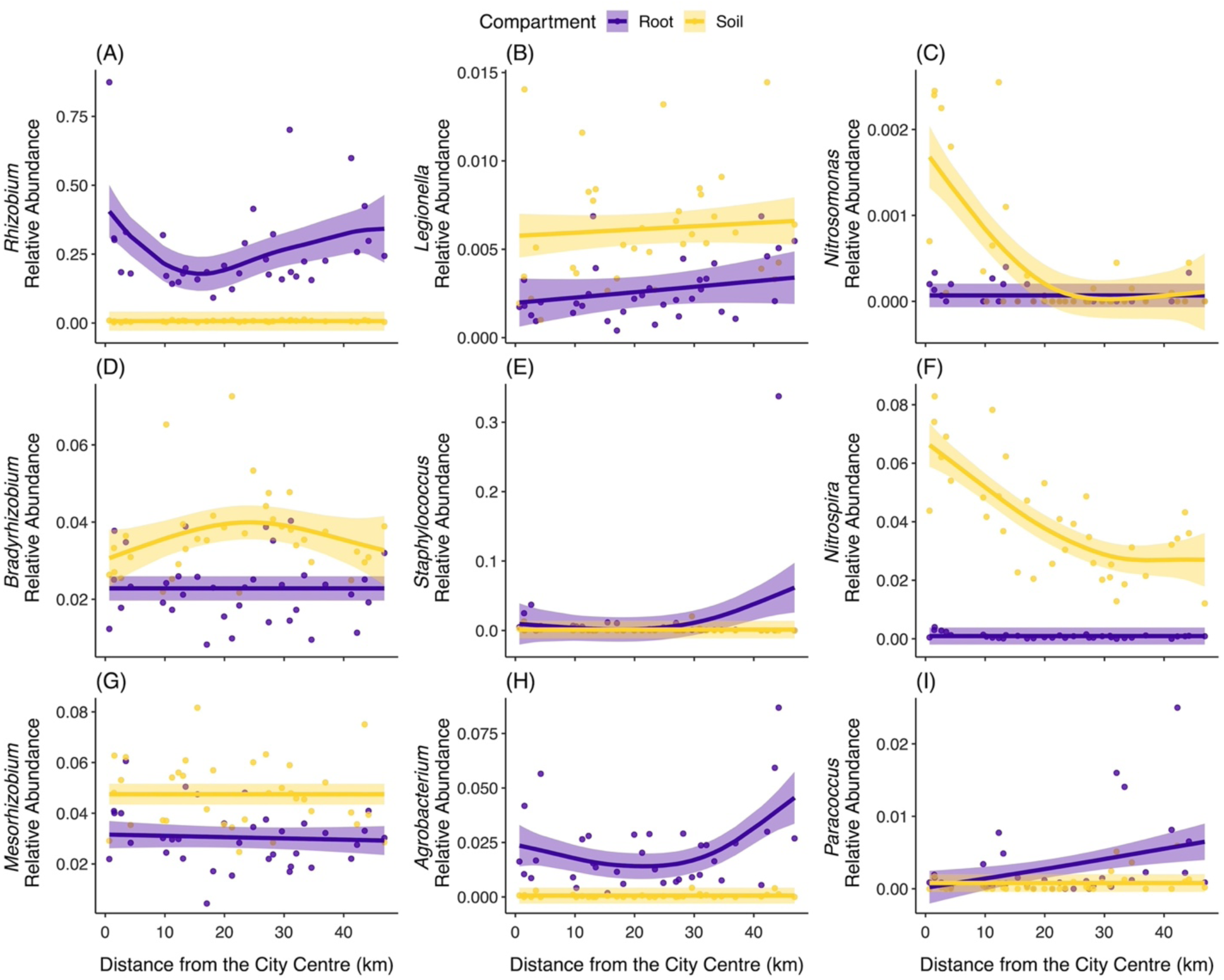
Relationships between the relative abundance of focal bacteria taxa and distance from the city centre. A subset of mutualistic (A, D, G), pathogenic (B, E, H), and ecosystem function (C, F, I) bacteria are presented. Points represent population means and lines are smoothed curves (± 95% confidence interval) from generalized additive models, with separate curves shown for the root and soil microbiome compartments (purple and yellow, respectively). Deviance explained by the models: *Rhizobium* (66.8%), *Legionella* (49.4%), *Nitrosomonas* (55.5%), *Bradyrhizobium* (39.7%), *Staphylococcus* (15.4%), *Nitrospira* (86.8%), *Mesorhizobium* (33.2%), *Agrobacterium* (56.4%), and *Paracoccus* (20.4%). Detailed statistics are provided in Tables S5-S7. Note: the scale of the y-axis varies by focal bacterium.

## Discussion

Our study demonstrates that urbanization influences the diversity and ecological functioning of soil and root microbiomes, but there were no direct or indirect effects of urbanization on plant microbiome community assembly. We found that soil alpha diversity was influenced by urbanization (Q1), with greater richness and phylogenetic diversity at the urban and rural limits of the gradient. Additionally, soil and root communities had distinct compositions, with greater variation in composition for root microbiomes compared to soil microbiomes (Q1). Urbanization directly and indirectly mediated soil microbiome assembly, particularly through soil C. Root microbiome assembly was not linked to urbanization, suggesting that the host plant acts as an additional filter on microbiome assembly independent of urbanization (Q2). Finally, we found differences in the abundance of bacteria with urbanization, including key mutualistic and pathogenic bacteria (Q3). Below, we discuss the broader ecological relevance and applied importance of these results.

### How does the diversity and composition of the microbiome vary with urbanization?

Root alpha diversity was not affected by urbanization, despite considerable non-linear variation in soil alpha diversity along the urbanization gradient. Previous research on microbes in urban environments has primarily focused soil communities [17–19,22–25,27], whereby the bacteria respond to the local environment and urban-driven changes to soil properties. In contrast, plant microbiome diversity is a function of local environmental conditions and the plant microbiome compartment [2–4]. Select studies of plant microbiome assembly in urban environments have primarily focused on bacterial communities on leaves (i.e., phyllosphere) [but see 29], with both positive [28] and negative [30] impacts of urbanization on bacterial microbiome assembly and diversity. As the phyllosphere is directly exposed to the environment, it could respond more strongly to environmental change [2–4,69]. Our study examined bacterial diversity in the highly plant-regulated endophytic root compartment [4,69], which, through the selective recruitment of bacteria from the soil, could buffer against changes in alpha diversity due to urbanization.

Urbanization did not act as a driver of biotic homogenization for either root or soil bacteria. Previous studies have found convergence in soil microbial community structure across sites categorized by human disturbance or in urban greenspaces [19,25,27]. Comparisons along an urban-rural gradient can yield a different pattern of diversity and composition than studies that compare among urban-rural pairs or categories (e.g., low, medium, high) of human disturbance. By using the gradient approach, we were able to detect non-linear variation in root beta dispersion and soil alpha diversity when categorical comparisons between urban and rural sites would have likely shown no effect of urbanization. Surprisingly, we also found that root bacterial microbiomes had greater variation in community composition than soil microbiomes. Greater beta diversity in the roots could result from population- or genotype-specific assembly [21,70,71]. Experiments manipulating soil properties, microbial inocula, and host population/genotype are needed to disentangle the extent of host control on and variation of root bacterial microbiome composition.

### What are the direct and indirect effects of urbanization on soil chemistry and microbiome assembly?

Root and soil microbiome assembly followed different pathways. Soil microbiome assembly was influenced by urbanization via soil total C. Carbon is an essential resource for free-living bacteria [72,73], as it can influence the abundance and biomass of microbes that can be supported, and urbanization reduced the amount of available soil total C (electronic supplementary material, figure S10). Root microbiome assembly was influenced by the soil microbiome composition, with cascading impacts on white clover δ^15^N. Urbanization also had indirect effects on white clover δ^15^N, whereby urbanization increased white clover total N (electronic supplementary material, figure S11). A previous study found that white clover δ^15^N was directly influenced by urbanization, with indirect links mediated through rhizobia and soil total N [31]. Pairing previous results with our current study, urbanization can directly alter total N and the source of N used by white clover (i.e., δ^15^N), with a potential role for the broader bacterial microbiome − in addition to *Rhizobium* − regulating N fixation. We did not find a direct or indirect link between urbanization root microbiome assembly, and we interpret this result as white clover acting as an additional filter to the soil compartment on root bacterial recruitment [2–4,69]. As root bacterial microbiomes varied in composition among populations, further work should investigate the ecosystem-level consequences of root microbiome variation and population-by-microbiome interactions on N fixation.

### How does the prevalence of mutualistic, pathogenic, and ecosystem function bacteria change with urbanization?

Bacteria comprising key functional groups differed with urbanization, with potential impacts for community interactions, ecosystem processes, and human and wildlife health. As the primary rhizobial symbiont of white clover, *Rhizobium* dominated the root compartment, peaking in abundance at the urban and rural limits of the gradient. Previous research showed that nodule density decreased with increasing urbanization [31], and rhizobia density is related to nodulation [74,75]. Relative abundances of *Rhizobium* could be maintained despite increased urbanization because it is not just the amount of *Rhizobium* that matters but having the ‘right’ *Rhizobium* that are locally-adapted to their plant hosts [76]. Local adaptation between white clover and *Rhizobium* has been documented in this system [76], and it could contribute to the observed patterns in *Rhizobium* relative abundance. In terms of important pathogenic bacteria, the plant pathogens *Agrobacterium* and *Rhizobacter* increased in rural and agricultural areas of the gradient [77] We also found that the human pathogens like *Legionella* [e.g., Legionnaires’ disease, pneumonia; 77,78] increased with decreasing urbanization, while *Clostridium* [e.g., cellulitis, botulism, tetanus; 79,80] were more abundant in urban areas. For each of the pathogenic bacteria, understanding their spatial distribution and abundance is important for management and public health because the soil and plant microbiomes can serve as a reservoir for transmission [26,82]. Our study was limited to 16S amplicon sequencing, but future work could use metagenomic approaches to identify species or strains of particular pathogenic interest. *Conclusions*

Our study provides compelling evidence that urbanization, a leading driver of environmental change, can affect the assembly and diversity of soil and root microbiomes. Urbanization did not decrease alpha diversity or homogenize community composition of soil microbiomes, in contrast to other studies [19,24,25,27], although soil bacterial diversity was lowest in suburban habitats. Root alpha diversity was not influenced by urbanization, but we unexpectedly found that root microbiomes exhibited greater variation in community composition than soil microbiomes and this variation was related to urbanization. We found indirect links between urbanization and soil microbiome assembly, but root microbiome assembly was independent of urbanization, which is likely due to control of the root compartment [4,69]. Given the links between soil microbiome composition and root microbiome assembly, observed variation in soil and root microbiome composition by population could have implications for plant-microbiome adaptation and associated community- and ecosystem-level consequences [83,84], although this requires further study and experimentation. Key mutualistic, pathogenic, and ecosystem functioning bacteria also varied with urbanization, with implications for nutrient cycling and plant and human pathogen management [10,11,26]. Given the extensive and increasing rate of urbanization [12–15], our study underscores the importance of examining how urban-driven environmental change alters the ecology and function of soil and root microbiomes [1,10,11,84].

## Ethics

This work did not require ethical approval from a human subject or animal welfare committee.

## Data Accessibility

Data, metadata, and R code are available on Zenodo (<u>10.5281/zenodo.11550118</u>). Sequences are available at NCBI under the BioProject accession <u>PRJNA1107685</u>.

## Declaration of AI Use

We have not used AI-assisted technologies in creating this article.

## Authors’ Contributions

**DMS:** conceptualization, investigation, methodology, formal analysis, visualization, data curation, writing-original draft, writing-review and editing, funding acquisition; **LR:** investigation, methodology, writing-review and editing; **MJ:** conceptualization, investigation, methodology, formal analysis, writing-review and editing, funding acquisition.

## Acknowledgements

We thank Lindsay Miles and Kelly Murray-Stoker for assisting with field work. Comments from Sophie Breitbart, Lucas Albano, Aude Caizergues, Ella Martin, and Nehal Naik greatly improved the manuscript. Megan Frederickson and John Stinchcombe provided guidance with study design and advice on statistical analyses.

