## Supplemental Methods for "Diversity and assembly of the microbiome of a leguminous plant along an urbanization gradient"

### Statistical Analyses

#### *Alpha Diversity*

Changes in alpha diversity with urbanization were evaluated using generalized additive mixed models [GAMMs; 1,2]. Generalized additive models are a flexible approach to analyzing nonlinear data, where nonparametric smoothing functions identify the relationship between the response and predictor [1–3].

We used four different metrics of alpha diversity: ASV richness, inverse Simpson, ASV evenness (inverse Simpson/ASV richness), and Faith's phylogenetic diversity (PD). Urbanization was quantified as distance from the city centre, HII, and mean ISC. Models were fitted as:

$$y_i \sim \text{intercept} + \text{Compartment} + \text{s}(x_i, \text{by} = \text{Compartment}) + \text{s}(\text{Population}) + e_i$$

where $y_i$ was the alpha diversity response, compartment (root and soil) was a parametric fixed effect, $x_i$ was a nonparametric predictor (distance from the city centre, HII, or mean ISC) that had separate smoothing functions for each compartment [indicated by s($x_i$, by = Compartment)], population was fitted as a nonparametric random intercept, and $e_i$ was the residual error associated with the model. Smooths for urbanization metrics were fitted using thin-plate regression splines with shrinkage terms [method call: bs = "ts" 4], and smooths for random intercepts were fitted using a ridge penalty [method call: bs = "re" 2]. All GAMMs were fitted using restricted maximum likelihood with smoothness selection [1,5,6].

To optimize smoothing and reduce overfitting, we manually set the basis size (i.e., how 'wiggly' the model can be) for the urbanization metrics (k = 10) and population (k = 35) and confirmed the basis size was appropriate using the `k.check()` function [7]. Moreover, we visually assessed the dispersion of residuals and homogeneity of variance for all GAMMs. High concurvity (i.e., collinearity) between variables precluded a single GAMM with all urbanization metrics.

*Beta Diversity and Community Composition*

We evaluated beta diversity as the separation between root and soil compartments, population, and their interaction using a principal coordinates analysis (PCoA) followed by a PERMANOVA [8,9]. We conducted separate PCoAs using Bray-Curtis, unweighted UniFrac, and weighted UniFrac dissimilarity matrices. Analyses for each of the dissimilarity matrices yielded similar results, so we focused on weighted UniFrac because: (1) it considers taxon abundances alongside phylogenetic diversity [10], and (2) it explained the greatest amount of variation across 2 PCoA axes (55.7%). We calculated each dissimilarity matrix on relative abundances, with dissimilarities compared using a PERMANOVA with 1000 permutations [`adonis2()` function; 10].

We further evaluated beta diversity by calculating the beta dispersion of community composition [11]. For each population, beta dispersion was calculated as the mean distance to the centroid in PCoA space, where the centroid represented the mean bacterial composition of all samples along the urbanization gradient. Beta dispersion was calculated using Bray-Curtis, unweighted UniFrac, and weighted UniFrac dissimilarity matrices, although we focus on the weighted UniFrac dissimilarity matrix. We compared beta dispersion by urbanization using generalized additive models [GAMs; 1]. We used the same model structure described above, except the nonparametric random intercept for population was dropped (i.e., only fixed effects included in the GAMs).

*Microbiome Assembly*

We used piecewise structural equation modeling (pSEM) to determine the hierarchical pathways of microbiome assembly [12,13]. By using this approach, we were able to test for direct and indirect effects of urbanization, soil nutrients, and soil microbial community composition and root microbiome assembly. Hypothesized pathways in pSEMs are specified based on prior knowledge [14–17], and any inference drawn from the pSEM depends on the ability of hypothesized causal relationships to be supported by the data [13,14].

We examined the overall fit of the hypothesized pSEM using Shipley's test of directed separation, which can indicate missing pathways between variables [18]. Our hypothesized pSEM was considered supported by the data when all plausible pathways were included [i.e., P-value for Fisher's C > 0.100; 14]. Results from the pSEM were reported as standardized path coefficients, which show the direction and magnitude of the relationship between variables and allow for comparisons within the pSEM [12,14–16,19].

*Differential Abundances*

We used the Analysis of Compositions of Microbiomes with Bias Correction (ANCOMBC) to detect differentially-abundant taxa in relation to urbanization [20]. In contrast to other differential abundance methods [e.g., DESeq2; 23] , ANCOMBC assumes that each sample is an unknown sampling fraction from its source ecosystem and this sampling fraction varies among samples [20,22]. ANCOMBC uses a linear regression framework with a sample-specific offset to correct for this bias.

We prepared data for analysis by using `phyloseq` to agglomerate the ASVs at the genus level. We then tested for differential abundance by fitting ANCOMBC with compartment (root and soil), an urbanization metric (distance, HII, or mean ISC), and their interaction as fixed terms. Taxa were only included if they were present at ≥ 10% of sites, and P-values were adjusted using the false discovery rate procedure [23]. Separate models were fitted for each urbanization metric due to collinearity.

We complemented ANCOMBC by evaluating how the relative abundances of focal bacterial taxa varied with urbanization. We identified putative mutualistic, pathogenic, and ecosystem functioning bacteria from the literature [26–29; table S1], and then searched our dataset for these specific bacteria. We calculated the relative abundance of each genus for each population, and then fitted GAMs of the form:

$$y_i \sim \text{intercept} + \text{Compartment} + \text{s}(x_i, \text{by} = \text{Compartment}) + \text{s(Population)} + e_i$$

where $y_i$ was the relative abundance of the focal bacterium, compartment (root and soil) was a parametric fixed effect, $x_i$ was a nonparametric predictor (distance from the city centre, HII, or mean ISC) that had separate smoothing functions for each compartment, and $e_i$ was the residual error associated with the model. Smooths for urbanization metrics were fitted using thin-plate regression splines with shrinkage terms [method call: bs = "ts" 4]. All GAMs were fitted using restricted maximum likelihood with smoothness selection [1,5,6]. We followed the same GAM evaluation procedure described above to check for overfitting, and high concurvity between variables precluded a single GAM for each bacterium with all urbanization metrics. We acknowledge this approach of targeted bacteria is not exhaustive, and we refer readers to the ANCOMBC results for a comprehensive analysis of the differential abundance of all bacteria.

*Soil and Leaf Nutrient and Isotope Analyses*

Changes in soil and leaf nutrients (% C and % N) and isotopes ($\delta^{13}$C and $\delta^{15}$N) were compared using GAMs. We fitted the GAMs as:

$$y_i = \text{intercept} + s(x_i) + e_i$$

where $y_i$ was the response (white clover $\delta^{15}N$, bulk soil N, or adjacent soil N), $x_i$ was the nonparametric predictor (distance from the urban center, HII, or mean ISC), and $e_i$ was the residual error. Thin-plate regression splines with shrinkage terms (method call: bs = "ts") were used to smooth all predictors [4]. All GAMs were fitted using restricted maximum likelihood with smoothness selection [1,5,6]. We followed the same GAM evaluation procedure described above to check for overfitting.

*Interpretation of Results*

We focused on estimates, variance explained, and effect sizes when interpreting statistical results [28–32]. For GAM(M)s, the effect size for the entire model is the percentage of deviance explained [1,2], which is analogous to unadjusted $R^2$. We quantified the effect size of individual model terms as partial eta-squared ($\eta^2_P$), which represents the proportion of variance explained by that term [33]; we quantified $\eta^2_P$ using the \`effectsize\` package [34]. For ANCOMBC, we emphasized the log-fold change in abundance of bacteria, although we filtered the dataset to focus on bacteria with $P < 0.100$. Similarly, we filtered the pSEM results to only include pathways with $P < 0.100$. Filtering the ANCOMBC and pSEM results was done for graphical clarity, and we provide full results for both ANCOMBC (electronic supplementary material, ANCOMBC Dataset) and the pSEM (electronic supplementary material, table S5).

*Statistical Software*

All of the above analyses were performed using R [version 4.2.3; 37] in the RStudio environment [version 2022.07.2; 38]. Data management and figure creation were facilitated using the \`tidyverse\` [37]and \`ggpubr\` [38] packages. Graphical and numerical evaluation of model fits was conducted using the \`performance\` package [39]. All data and R code will be deposited on Zenodo, and sequences will be available at NCBI under a BioProject accession.
