## Supplemental Tables and Figures for "Diversity and assembly of the microbiome of a leguminous plant along an urbanization gradient"

### Urban Microbiome – Supplemental Tables and Figures

#### Supplemental Tables

**Table S1:** List of focal bacteria related to mutualistic or pathogenic roles and broader ecosystem functioning, with a brief description of each focal bacterium in relation to its functional group.

| Functional Group | Genus | Brief Description | Citation(s) |
|---|---|---|---|
| Mutualistic | *Allorhizobium* | Symbiotic N-fixation | [1–4] |
| | *Bradyrhizobium* | Symbiotic N-fixation | [1–4] |
| | *Mesorhizobium* | Symbiotic N-fixation | [1–4] |
| | *Pararhizobium* | Symbiotic N-fixation | [2–4] |
| | *Rhizobium* | Symbiotic N-fixation | [1–4] |
| | *Shinella* | Symbiotic N-fixation | [3] |
| Pathogenic | *Acinetobacter* | Human pathogen that can cause illness such as pneumonia, bacteraemia, meningitis, and pulmonary infections; some strains have developed antibiotic resistance | [5,6] |
| | *Agrobacterium* | Plant pathogen (e.g., crown gall disease) | [7–9] |
| | *Clostridium* | Human pathogen that can cause illnesses like cellulitis, botulism, and tetanus | [7,10] |
| | *Legionella* | Human pathogen, particularly for immunosuppressed people; examples of illnesses include pneumonia, Legionnaires' disease, and extrapulmonary infection | [11,12] |
| | *Mycobacterium* | Infect mammalian hosts; certain species can cause illnesses like tuberculosis, leprosy, chronic pulmonary disease, and skin infections | [13,14] |
| | *Paenibacillus* | Beneficial to plants by producing antimicrobial compounds that suppress pathogens; considered pathogenic because it suppresses other pathogens and is not a mutualistic interaction *stricto sensu* | [1,15] |
| | *Rhizobacter* | Plant pathogen (e.g., gall inducing) | [7,16] |
| | *Sphingomonas* | Beneficial to plants by producing antimicrobial compounds that suppress pathogens; considered pathogenic because it suppresses other pathogens and is not a mutualistic interaction *stricto sensu* | [17,18] |
| | *Staphylococcus* | Pathogen of human and other mammals (e.g. staph infections); some strains have developed antibiotic resistance | [19,20] |
| | *Stenotrophomonas* | Opportunistic pathogen of plants (*S. maltophilia* is the only species known cause human disease); some species can be beneficial to plant growth and ecosystem processes, but *S. maltophilia* is most common | [17,21] |
| | *Williamsia* | Human pathogen that can cause illnesses like pulmonary infection, bacteraemia, and endophthalmitis | [22–24] |
| | *Xanthomonas* | Plant pathogen, particularly for crop plant species | [7,8] |
| Ecosystem Functioning | *Aciditerrimonas* | Iron reduction | [25,26] |
| | *Cytophaga* | Decomposition of cellulose | [27] |
| | *Desulfocapsa* | Sulfur cycling, specifically the reduction of sulfur | [28] |
| | *Desulfosporosinus* | Sulfur cycling, specifically the reduction of sulfur | [29] |
| | *Desulfuromonas* | Sulfur cycling, specifically the reduction of S to $H_2S$ | [30,31] |
| | *Devosia* | Bioremediation | [32,33] |

| Table S1 *(continued)* | | | |
|---|---|---|---|
| Ecosystem Functioning | *Nitrosomonas* | Nitrogen cycling, oxidation of $NH_3$ | [34] |
| | *Nitrosospira* | Nitrogen cycling, oxidation of $NH_3$ | [34] |
| | *Nitrospira* | Nitrogen cycling, oxidation of $NO_2^-$ | [34] |
| | *Novosphingobium* | Degradation of aromatic compounds | [35,36] |
| | *Paracoccus* | Nitrogen cycling, specifically the reduction of $NO_3^-$ to $NO_2^-$, NO, $N_20$, and $N_2$ | [37] |
| | *Phyllobacterium* | Promotes plant growth and tolerance to environmental stress (e.g., drought) | [38,39] |
| | *Sphingobium* | Biodegradation of environmental contaminants | [40] |
| | *Sphingopyxis* | Biodegradation and remediation of environmental contaminants | [41] |
| | *Streptomyces* | Plant growth promotion; source of antibiotics | [42,43] |

**Table S2:** Summary of the alpha diversity-by-urbanization generalized additive mixed models. We provide the P-value and effect size (partial eta-squared, $\eta^2_P$) for each model term for each measure of alpha diversity [richness, inverse Simpson, evenness, and Faith's phylogenetic diversity (PD)]. Compartment is the parametric term for microbiome compartment (root or soil), with Urbanization:Root and Urbanization:Soil reflecting the smoothed term for the respective urbanization metric for each compartment. Population is the random intercept of population identity.

| Urbanization Metric | Compartment | Urbanization:Root | Urbanization:Soil | Population |
|---|---|---|---|---|
| **Richness** | | | | |
| Distance from the City Centre | $P < 0.001$, $\eta^2_P = 0.672$ | $P = 0.194$, $\eta^2_P < 0.001$ | $P < 0.001$, $\eta^2_P = 0.026$ | $P = 0.123$, $\eta^2_P = 0.007$ |
| Human Influence Index | $P < 0.001$, $\eta^2_P = 0.673$ | $P = 0.156$, $\eta^2_P < 0.001$ | $P = 0.002$, $\eta^2_P = 0.078$ | $P = 0.068$, $\eta^2_P < 0.001$ |
| Mean Impervious Surface Cover | $P < 0.001$, $\eta^2_P = 0.667$ | $P = 0.138$, $\eta^2_P < 0.001$ | $P < 0.001$, $\eta^2_P = 0.004$ | $P = 0.437$, $\eta^2_P = 0.013$ |
| **Inverse Simpson** | | | | |
| Distance from the City Centre | $P < 0.001$, $\eta^2_P = 0.816$ | $P = 0.845$, $\eta^2_P \cong 0$ | $P < 0.001$, $\eta^2_P = 0.129$ | $P < 0.001$, $\eta^2_P = 0.247$ |
| Human Influence Index | $P < 0.001$, $\eta^2_P = 0.815$ | $P = 0.989$, $\eta^2_P \cong 0$ | $P < 0.001$, $\eta^2_P = 0.132$ | $P < 0.001$, $\eta^2_P = 0.235$ |
| Mean Impervious Surface Cover | $P < 0.001$, $\eta^2_P = 0.818$ | $P = 0.565$, $\eta^2_P \cong 0$ | $P < 0.001$, $\eta^2_P = 0.196$ | $P < 0.001$, $\eta^2_P = 0.196$ |
| **Evenness** | | | | |
| Distance from the City Centre | $P < 0.001$, $\eta^2_P = 0.610$ | $P = 0.552$, $\eta^2_P \cong 0$ | $P = 0.377$, $\eta^2_P < 0.001$ | $P < 0.001$, $\eta^2_P = 0.186$ |
| Human Influence Index | $P < 0.001$, $\eta^2_P = 0.610$ | $P = 0.309$, $\eta^2_P \cong 0$ | $P = 0.443$, $\eta^2_P < 0.001$ | $P < 0.001$, $\eta^2_P = 0.183$ |
| Mean Impervious Surface Cover | $P < 0.001$, $\eta^2_P = 0.676$ | $P = 0.859$, $\eta^2_P \cong 0$ | $P = 0.193$, $\eta^2_P < 0.001$ | $P < 0.001$, $\eta^2_P = 0.186$ |
| **Faith's PD** | | | | |
| Distance from the City Centre | $P < 0.001$, $\eta^2_P = 0.719$ | $P = 0.147$, $\eta^2_P < 0.001$ | $P < 0.001$, $\eta^2_P = 0.053$ | $P = 0.079$, $\eta^2_P = 0.011$ |
| Human Influence Index | $P < 0.001$, $\eta^2_P = 0.727$ | $P = 0.079$, $\eta^2_P < 0.001$ | $P < 0.001$, $\eta^2_P = 0.118$ | $P = 0.022$, $\eta^2_P < 0.001$ |
| Mean Impervious Surface Cover | $P < 0.001$, $\eta^2_P = 0.719$ | $P = 0.084$, $\eta^2_P = 0.001$ | $P < 0.001$, $\eta^2_P = 0.007$ | $P = 0.382$, $\eta^2_P = 0.023$ |

**Table S3:** Summary of the beta dispersion-by-urbanization generalized additive models. We fitted separate models for each urbanization metric and distance matrix (Bray-Curtis, UniFrac, and weighted UniFrac). We provide the P-value and effect size (partial eta-squared, $\eta^2_P$) for each model term. Compartment is the parametric term for microbiome compartment (root or soil), with Urbanization:Root and Urbanization:Soil reflecting the smoothed term for the respective urbanization metric for each compartment.

| Urbanization Metric | Compartment | Urbanization:Root | Urbanization:Soil |
|---|---|---|---|
| **Bray-Curtis** | | | |
| Distance from the City Centre | $P < 0.001$, $\eta^2_P = 0.625$ | $P = 0.012$, $\eta^2_P = 0.033$ | $P = 0.866$, $\eta^2_P < 0.001$ |
| Human Influence Index | $P < 0.001$, $\eta^2_P = 0.588$ | $P = 0.288$, $\eta^2_P < 0.001$ | $P = 0.367$, $\eta^2_P < 0.001$ |
| Mean Impervious Surface Cover | $P < 0.001$, $\eta^2_P = 0.587$ | $P = 0.367$, $\eta^2_P < 0.001$ | $P = 0.508$, $\eta^2_P < 0.001$ |
| **UniFrac** | | | |
| Distance from the City Centre | $P < 0.001$, $\eta^2_P = 0.502$ | $P = 0.921$, $\eta^2_P < 0.001$ | $P = 0.319$, $\eta^2_P < 0.001$ |
| Human Influence Index | $P < 0.001$, $\eta^2_P = 0.516$ | $P = 0.306$, $\eta^2_P < 0.001$ | $P = 0.044$, $\eta^2_P = 0.004$ |
| Mean Impervious Surface Cover | $P < 0.001$, $\eta^2_P = 0.509$ | $P = 0.373$, $\eta^2_P < 0.001$ | $P = 0.015$, $\eta^2_P = 0.029$ |
| **Weighted UniFrac** | | | |
| Distance from the City Centre | $P < 0.001$, $\eta^2_P = 0.651$ | $P = 0.008$, $\eta^2_P = 0.037$ | $P = 0.987$, $\eta^2_P < 0.001$ |
| Human Influence Index | $P < 0.001$, $\eta^2_P = 0.611$ | $P = 0.355$, $\eta^2_P < 0.001$ | $P = 0.614$, $\eta^2_P < 0.001$ |
| Mean Impervious Surface Cover | $P < 0.001$, $\eta^2_P = 0.611$ | $P = 0.471$, $\eta^2_P < 0.001$ | $P = 0.582$, $\eta^2_P < 0.001$ |

**Table S4:** Path coefficients for each pathway in the piecewise structural equation model (pSEM). We provide each structural equation in the pSEM, with causal pathways indicated by ~ and correlational pathways indicated by ~~. For each structural equation, we report the unstandardized path coefficient (Estimate), standard error of the unstandardized, path coefficient (SE), standardized path coefficient (Path Coef), z-statistic, and P-value. Variables are abbreviated as: Distance = distance from the city centre, HII = Human Influence Index, ISC = impervious surface cover, % C = total carbon, % N = total nitrogen, Soil PCoA 1= soil microbiome principal coordinates analysis axis 1, Soil PCoA 2= soil microbiome principal coordinates analysis axis 2, Root PCoA 1= root microbiome principal coordinates analysis axis 1, and Root PCoA 2= root microbiome principal coordinates analysis axis 2.

| Structural Equation | Estimate | SE | Path Coefficient | P-value |
|---|---|---|---|---|
| Mean ISC ~ Distance | -1.491 | 0.202 | -0.790 | < 0.001 |
| HII ~ Distance | -0.528 | 0.071 | -0.792 | < 0.001 |
| Mean ISC ~~ HII | 0.240 | NA | 0.240 | 0.085 |
| Soil % C ~ Mean ISC | 0.019 | 0.015 | 0.287 | 0.209 |
| Soil % C ~ HII | -0.115 | 0.041 | -0.626 | 0.009 |
| Soil $\delta^{13}C$ ~ Distance* | 0.243 | 0.104 | 0.638 | 0.026 |
| Soil $\delta^{13}C$ ~ Mean ISC | 0.010 | 0.048 | 0.049 | 0.839 |
| Soil $\delta^{13}C$ ~ HII | -0.011 | 0.136 | -0.019 | 0.938 |
| Soil % C ~~ Soil $\delta^{13}C$ | 0.398 | NA | 0.398 | 0.010 |
| Soil % C ~~ Soil % N | 0.366 | NA | 0.366 | 0.017 |
| Soil % N ~ Distance* | -0.005 | 0.002 | -0.686 | 0.034 |
| Soil % N ~ Mean ISC | < 0.001 | 0.001 | -0.070 | 0.799 |
| Soil % N ~ HII | -0.003 | 0.003 | -0.304 | 0.272 |
| Soil $\delta^{15}N$ ~ Mean ISC | 0.012 | 0.011 | 0.237 | 0.260 |
| Soil $\delta^{15}N$ ~ HII | 0.056 | 0.030 | 0.386 | 0.071 |
| Soil % N ~~ Soil $\delta^{15}N$ | 0.349 | NA | 0.349 | 0.021 |
| Soil % N ~~ Soil $\delta^{13}C$* | -0.475 | NA | -0.475 | 0.002 |
| Leaf % C ~ Mean ISC | 0.023 | 0.013 | 0.413 | 0.091 |
| Leaf % C ~ HII | -0.077 | 0.041 | -0.487 | 0.069 |
| Leaf % C ~ Soil $\delta^{15}N$* | 0.462 | 0.215 | 0.426 | 0.040 |
| Leaf % C ~ Root PCoA 1 | 5.762 | 14.113 | 0.071 | 0.686 |
| Leaf % C ~ Root PCoA 2 | 0.025 | 7.352 | 0.001 | 0.997 |
| Leaf $\delta^{13}C$ ~ Mean ISC | -0.008 | 0.008 | -0.229 | 0.341 |
| Leaf $\delta^{13}C$ ~ HII | 0.003 | 0.024 | 0.027 | 0.916 |
| Leaf $\delta^{13}C$ ~ Root PCoA 1 | -19.326 | 8.723 | -0.390 | 0.034 |
| Leaf $\delta^{13}C$ ~ Root PCoA 2 | 5.302 | 4.554 | 0.221 | 0.254 |
| Leaf % C ~~ Leaf $\delta^{13}C$ | 0.031 | NA | 0.031 | 0.430 |
| Leaf % C ~~ Leaf % N | 0.421 | NA | 0.421 | 0.006 |
| Leaf % N ~ Mean ISC | 0.007 | 0.004 | 0.361 | 0.104 |
| Leaf % N ~ HII | -0.018 | 0.013 | -0.322 | 0.179 |
| Leaf % N ~ Soil % N | 0.247 | 0.949 | 0.045 | 0.797 |
| Leaf % N ~ Soil $\delta^{15}N$ | 0.211 | 0.075 | 0.544 | 0.009 |
| Leaf % N ~ Root PCoA 1 | 3.640 | 4.805 | 0.125 | 0.455 |
| Leaf % N ~ Root PCoA 2 | 1.822 | 2.380 | 0.129 | 0.450 |
| Leaf $\delta^{15}N$ ~ Mean ISC | 0.004 | 0.005 | 0.209 | 0.400 |
| Leaf $\delta^{15}N$ ~ HII | -0.013 | 0.015 | -0.230 | 0.395 |
| Leaf $\delta^{15}N$ ~ Soil % N | 0.547 | 1.057 | 0.102 | 0.609 |
| Leaf $\delta^{15}N$ ~ Soil $\delta^{15}N$ | 0.088 | 0.084 | 0.233 | 0.302 |
| Leaf $\delta^{15}N$ ~ Root PCoA 1 | 7.547 | 5.351 | 0.265 | 0.170 |
| Leaf $\delta^{15}N$ ~ Root PCoA 2 | -4.976 | 2.650 | -0.361 | 0.071 |
| Leaf % N ~~ Leaf $\delta^{15}N$ | 0.270 | NA | 0.270 | 0.061 |
| Soil PCoA 1 ~ Mean ISC | < 0.001 | < 0.001 | -0.409 | 0.146 |
| Soil PCoA 1 ~ HII | 0.001 | < 0.001 | 0.363 | 0.216 |
| Soil PCoA 1 ~ Soil % C | 0.003 | 0.004 | 0.222 | 0.479 |
| Soil PCoA 1 ~ Soil $\delta^{13}C$ | < 0.001 | 0.001 | -0.083 | 0.810 |
| Soil PCoA 1 ~ Soil % N | 0.015 | 0.076 | 0.070 | 0.844 |
| Soil PCoA 1 ~ Soil $\delta^{15}N$ | 0.004 | 0.004 | 0.273 | 0.267 |
| Soil PCoA 2 ~ Mean ISC | < 0.001 | < 0.001 | -0.099 | 0.598 |
| Soil PCoA 2 ~ HII | < 0.001 | < 0.001 | -0.032 | 0.869 |
| Soil PCoA 2 ~ Soil % C | 0.010 | 0.003 | 0.813 | 0.001 |
| Soil PCoA 2 ~ Soil $\delta^{13}C$ | -0.001 | < 0.001 | -0.318 | 0.182 |
| Soil PCoA 2 ~ Soil % N | -0.044 | 0.052 | -0.205 | 0.398 |
| Soil PCoA 2 ~ Soil $\delta^{15}N$ | -0.004 | 0.003 | -0.251 | 0.137 |
| Soil PCoA 1 ~~ Soil PCoA 2 | -0.108 | NA | -0.108 | 0.271 |

**Table S4** ***(continued)***

| | | | | |
|---|---|---|---|---|
| Root PCoA 1 ~ Mean ISC | < 0.001 | < 0.001 | 0.043 | 0.872 |
| Root PCoA 1 ~ HII | < 0.001 | < 0.001 | 0.077 | 0.781 |
| Root PCoA 1 ~ Soil PCoA 1 | 0.383 | 0.153 | 0.442 | 0.019 |
| Root PCoA 1 ~ Soil PCoA 2 | 0.002 | 0.226 | 0.003 | 0.991 |
| Root PCoA 1 ~ Soil % C | < 0.001 | 0.004 | 0.028 | 0.939 |
| Root PCoA 1 ~ Soil $\delta^{13}C$ | 0.001 | 0.001 | 0.200 | 0.549 |
| Root PCoA 1 ~ Soil % N | 0.076 | 0.062 | 0.405 | 0.232 |
| Root PCoA 1 ~ Soil $\delta^{15}N$ | -0.005 | 0.003 | -0.370 | 0.130 |
| Root PCoA 2 ~ Soil PCoA 1 | 0.039 | 0.313 | 0.022 | 0.901 |
| Root PCoA 2 ~ Soil PCoA 2 | 0.653 | 0.463 | 0.363 | 0.179 |
| Root PCoA 2 ~ Soil % C | 0.004 | 0.008 | 0.181 | 0.618 |
| Root PCoA 2 ~ Soil $\delta^{13}C$ | < 0.001 | 0.002 | 0.060 | 0.856 |
| Root PCoA 2 ~ Soil % N | 0.017 | 0.127 | 0.043 | 0.896 |
| Root PCoA 2 ~ Soil $\delta^{15}N$ | 0.002 | 0.007 | 0.091 | 0.703 |
| Root PCoA 2 ~ Mean ISC | < 0.001 | < 0.001 | 0.005 | 0.985 |
| Root PCoA 2 ~ HII | -0.001 | 0.001 | -0.168 | 0.541 |
| Root PCoA 1 ~~ Root PCoA 2 | 0.294 | NA | 0.294 | 0.046 |

**Note:** Pathways added to the pSEM following tests of directed separation are indicated by *

**Table S5:** Summary of the mutualistic bacteria relative abundance generalized additive models. We provide the P-value and effect size (partial eta-squared, $\eta^2_P$) for each model term. Compartment is the parametric term for microbiome compartment (root or soil), with Urbanization:Root and Urbanization:Soil reflecting the smoothed term for the respective urbanization metric for each compartment.

| Urbanization Metric | Compartment | Urbanization:Root | Urbanization:Soil |
|---|---|---|---|
| ***Allorhizobium*** | | | |
| Distance from the City Centre | $P < 0.001$, $\eta^2_P = 0.163$ | $P < 0.001$, $\eta^2_P = 0.093$ | $P \cong 1.000$, $\eta^2_P < 0.001$ |
| Human Influence Index | $P = 0.002$, $\eta^2_P = 0.139$ | $P = 0.002$, $\eta^2_P = 0.014$ | $P = 0.997$, $\eta^2_P < 0.001$ |
| Mean Impervious Surface Cover | $P = 0.003$, $\eta^2_P = 0.127$ | $P = 0.093$, $\eta^2_P = 0.002$ | $P \cong 1.000$, $\eta^2_P < 0.001$ |
| ***Bradyrhizobium*** | | | |
| Distance from the City Centre | $P < 0.001$, $\eta^2_P = 0.355$ | $P = 0.720$, $\eta^2_P < 0.001$ | $P = 0.055$, $\eta^2_P = 0.017$ |
| Human Influence Index | $P < 0.001$, $\eta^2_P = 0.332$ | $P = 0.675$, $\eta^2_P < 0.001$ | $P = 0.413$, $\eta^2_P < 0.001$ |
| Mean Impervious Surface Cover | $P < 0.001$, $\eta^2_P = 0.357$ | $P = 0.949$, $\eta^2_P < 0.001$ | $P = 0.048$, $\eta^2_P = 0.018$ |
| ***Mesorhizobium*** | | | |
| Distance from the City Centre | $P < 0.001$, $\eta^2_P = 0.328$ | $P = 0.244$, $\eta^2_P < 0.001$ | $P = 0.443$, $\eta^2_P < 0.001$ |
| Human Influence Index | $P < 0.001$, $\eta^2_P = 0.360$ | $P = 0.311$, $\eta^2_P < 0.001$ | $P = 0.016$, $\eta^2_P = 0.031$ |
| Mean Impervious Surface Cover | $P < 0.001$, $\eta^2_P = 0.331$ | $P = 0.163$, $\eta^2_P = 0.001$ | $P = 0.713$, $\eta^2_P < 0.001$ |
| ***Pararhizobium*** | | | |
| Distance from the City Centre | $P < 0.001$, $\eta^2_P = 0.679$ | $P = 0.116$, $\eta^2_P = 0.002$ | $P = 0.785$, $\eta^2_P < 0.001$ |
| Human Influence Index | $P < 0.001$, $\eta^2_P = 0.676$ | $P = 0.192$, $\eta^2_P = 0.001$ | $P = 0.896$, $\eta^2_P < 0.001$ |
| Mean Impervious Surface Cover | $P < 0.001$, $\eta^2_P = 0.675$ | $P = 0.219$, $\eta^2_P < 0.001$ | $P = 0.853$, $\eta^2_P < 0.001$ |
| ***Rhizobium*** | | | |
| Distance from the City Centre | $P < 0.001$, $\eta^2_P = 0.628$ | $P = 0.001$, $\eta^2_P = 0.088$ | $P = 0.995$, $\eta^2_P < 0.001$ |
| Human Influence Index | $P < 0.001$, $\eta^2_P = 0.568$ | $P = 0.119$, $\eta^2_P = 0.001$ | $P = 0.967$, $\eta^2_P < 0.001$ |
| Mean Impervious Surface Cover | $P < 0.001$, $\eta^2_P = 0.701$ | $P < 0.001$, $\eta^2_P = 0.329$ | $P = 0.997$, $\eta^2_P < 0.001$ |
| ***Shinella*** | | | |
| Distance from the City Centre | $P < 0.001$, $\eta^2_P = 0.375$ | $P = 0.016$, $\eta^2_P = 0.007$ | $P = 0.997$, $\eta^2_P < 0.001$ |
| Human Influence Index | $P < 0.001$, $\eta^2_P = 0.390$ | $P = 0.001$, $\eta^2_P = 0.015$ | $P \cong 1.000$, $\eta^2_P < 0.001$ |
| Mean Impervious Surface Cover | $P < 0.001$, $\eta^2_P = 0.364$ | $P = 0.078$, $\eta^2_P = 0.003$ | $P = 0.996$, $\eta^2_P < 0.001$ |

**Table S6:** Summary of the pathogenic bacteria relative abundance generalized additive models. We provide the P-value and effect size (partial eta-squared, $\eta^2_P$) for each model term. Compartment is the parametric term for microbiome compartment (root or soil), with Urbanization:Root and Urbanization:Soil reflecting the smoothed term for the respective urbanization metric for each compartment.

| Urbanization Metric | Compartment | Urbanization:Root | Urbanization:Soil |
|---|---|---|---|
| ***Acinetobacter*** | | | |
| Distance from the City Centre | $P = 0.116, \eta^2_P = 0.036$ | $P = 0.015, \eta^2_P = 0.007$ | $P = 0.376, \eta^2_P < 0.001$ |
| Human Influence Index | $P = 0.127, \eta^2_P = 0.034$ | $P = 0.209, \eta^2_P < 0.001$ | $P = 0.385, \eta^2_P < 0.001$ |
| Mean Impervious Surface Cover | $P = 0.120, \eta^2_P = 0.036$ | $P = 0.042, \eta^2_P = 0.004$ | $P = 0.655, \eta^2_P < 0.001$ |
| ***Agrobacterium*** | | | |
| Distance from the City Centre | $P < 0.001, \eta^2_P = 0.478$ | $P < 0.001, \eta^2_P = 0.090$ | $P = 0.934, \eta^2_P < 0.001$ |
| Human Influence Index | $P < 0.001, \eta^2_P = 0.586$ | $P < 0.001, \eta^2_P = 0.360$ | $P = 0.845, \eta^2_P < 0.001$ |
| Mean Impervious Surface Cover | $P < 0.001, \eta^2_P = 0.444$ | $P = 0.006, \eta^2_P = 0.040$ | $P \cong 1.000, \eta^2_P < 0.001$ |
| ***Clostridium*** | | | |
| Distance from the City Centre | $P < 0.001, \eta^2_P = 0.507$ | $P = 0.793, \eta^2_P < 0.001$ | $P = 0.006, \eta^2_P = 0.010$ |
| Human Influence Index | $P < 0.001, \eta^2_P = 0.484$ | $P = 0.786, \eta^2_P < 0.001$ | $P = 0.165, \eta^2_P = 0.001$ |
| Mean Impervious Surface Cover | $P < 0.001, \eta^2_P = 0.499$ | $P = 0.655, \eta^2_P < 0.001$ | $P = 0.020, \eta^2_P = 0.007$ |
| ***Legionella*** | | | |
| Distance from the City Centre | $P < 0.001, \eta^2_P = 0.327$ | $P = 0.119, \eta^2_P = 0.001$ | $P = 0.192, \eta^2_P = 0.001$ |
| Human Influence Index | $P < 0.001, \eta^2_P = 0.319$ | $P = 0.232, \eta^2_P < 0.001$ | $P = 0.587, \eta^2_P < 0.001$ |
| Mean Impervious Surface Cover | $P < 0.001, \eta^2_P = 0.330$ | $P = 0.139, \eta^2_P = 0.001$ | $P = 0.111, \eta^2_P = 0.002$ |
| ***Mycobacterium*** | | | |
| Distance from the City Centre | $P < 0.001, \eta^2_P = 0.597$ | $P = 0.644, \eta^2_P < 0.001$ | $P = 0.153, \eta^2_P = 0.001$ |
| Human Influence Index | $P < 0.001, \eta^2_P = 0.595$ | $P = 0.221, \eta^2_P < 0.001$ | $P = 0.295, \eta^2_P < 0.001$ |
| Mean Impervious Surface Cover | $P < 0.001, \eta^2_P = 0.592$ | $P = 0.551, \eta^2_P < 0.001$ | $P = 0.304, \eta^2_P < 0.001$ |
| ***Paenibacillus*** | | | |
| Distance from the City Centre | $P < 0.001, \eta^2_P = 0.372$ | $P = 0.614, \eta^2_P < 0.001$ | $P = 0.008, \eta^2_P = 0.010$ |
| Human Influence Index | $P < 0.001, \eta^2_P = 0.371$ | $P = 0.111, \eta^2_P = 0.002$ | $P = 0.031, \eta^2_P = 0.005$ |
| Mean Impervious Surface Cover | $P < 0.001, \eta^2_P = 0.354$ | $P = 0.619, \eta^2_P < 0.001$ | $P = 0.154, \eta^2_P = 0.001$ |
| ***Rhizobacter*** | | | |
| Distance from the City Centre | $P < 0.001, \eta^2_P = 0.642$ | $P = 0.003, \eta^2_P = 0.083$ | $P = 0.728, \eta^2_P < 0.001$ |
| Human Influence Index | $P < 0.001, \eta^2_P = 0.592$ | $P = 0.078, \eta^2_P = 0.003$ | $P = 0.786, \eta^2_P < 0.001$ |
| Mean Impervious Surface Cover | $P < 0.001, \eta^2_P = 0.591$ | $P = 0.094, \eta^2_P = 0.002$ | $P = 0.793, \eta^2_P < 0.001$ |
| ***Sphingomonas*** | | | |
| Distance from the City Centre | $P < 0.001, \eta^2_P = 0.538$ | $P < 0.001, \eta^2_P = 0.081$ | $P = 0.405, \eta^2_P < 0.001$ |
| Human Influence Index | $P < 0.001, \eta^2_P = 0.461$ | $P = 0.885, \eta^2_P < 0.001$ | $P = 0.998, \eta^2_P < 0.001$ |
| Mean Impervious Surface Cover | $P < 0.001, \eta^2_P = 0.461$ | $P = 0.751, \eta^2_P < 0.001$ | $P = 0.774, \eta^2_P < 0.001$ |
| ***Staphylococcus*** | | | |
| Distance from the City Centre | $P = 0.180, \eta^2_P = 0.027$ | $P = 0.012, \eta^2_P = 0.026$ | $P = 0.956, \eta^2_P < 0.001$ |
| Human Influence Index | $P = 0.104, \eta^2_P = 0.040$ | $P < 0.001, \eta^2_P = 0.214$ | $P = 0.921, \eta^2_P = 0.031$ |
| Mean Impervious Surface Cover | $P = 0.201, \eta^2_P = 0.024$ | $P = 0.128, \eta^2_P = 0.001$ | $P = 0.822, \eta^2_P < 0.001$ |
| ***Stenotrophomonas*** | | | |
| Distance from the City Centre | $P < 0.001, \eta^2_P = 0.240$ | $P = 0.035, \eta^2_P = 0.005$ | $P \cong 1.000, \eta^2_P < 0.001$ |
| Human Influence Index | $P < 0.001, \eta^2_P = 0.231$ | $P = 0.199, \eta^2_P = 0.001$ | $P = 0.862, \eta^2_P < 0.001$ |
| Mean Impervious Surface Cover | $P < 0.001, \eta^2_P = 0.236$ | $P = 0.077, \eta^2_P = 0.003$ | $P = 0.994, \eta^2_P < 0.001$ |
| ***Williamsia*** | | | |
| Distance from the City Centre | $P = 0.342, \eta^2_P = 0.013$ | $P = 0.772, \eta^2_P < 0.001$ | $P = 0.055, \eta^2_P = 0.003$ |
| Human Influence Index | $P = 0.341, \eta^2_P = 0.014$ | $P = 0.989, \eta^2_P < 0.001$ | $P = 0.043, \eta^2_P = 0.004$ |
| Mean Impervious Surface Cover | $P = 0.346, \eta^2_P = 0.013$ | $P = 0.797, \eta^2_P < 0.001$ | $P = 0.103, \eta^2_P = 0.002$ |
| ***Xanthomonas*** | | | |
| Distance from the City Centre | $P < 0.001, \eta^2_P = 0.213$ | $P = 0.011, \eta^2_P = 0.009$ | $P = 0.922, \eta^2_P < 0.001$ |
| Human Influence Index | $P < 0.001, \eta^2_P = 0.226$ | $P = 0.001, \eta^2_P = 0.018$ | $P = 0.794, \eta^2_P < 0.001$ |
| Mean Impervious Surface Cover | $P < 0.001, \eta^2_P = 0.210$ | $P = 0.025, \eta^2_P = 0.006$ | $P = 0.997, \eta^2_P < 0.001$ |

**Table S7:** Summary of the ecosystem function bacteria relative abundance generalized additive models. We provide the P-value and effect size (partial eta-squared, $\eta^2_P$) for each model term. Compartment is the parametric term for microbiome compartment (root or soil), with Urbanization:Root and Urbanization:Soil reflecting the smoothed term for the respective urbanization metric for each compartment.

| Urbanization Metric | Compartment | Urbanization:Root | Urbanization:Soil |
|---|---|---|---|
| ***Aciditerrimonas*** | | | |
| Distance from the City Centre | P < 0.001, $\eta^2_P$ = 0.387 | P = 0.989, $\eta^2_P$ < 0.001 | P < 0.001, $\eta^2_P$ = 0.066 |
| Human Influence Index | P < 0.001, $\eta^2_P$ = 0.327 | P = 0.885, $\eta^2_P$ < 0.001 | P < 0.001, $\eta^2_P$ = 0.020 |
| Mean Impervious Surface Cover | P < 0.001, $\eta^2_P$ = 0.315 | P = 0.860, $\eta^2_P$ < 0.001 | P < 0.001, $\eta^2_P$ = 0.013 |
| ***Cytophaga*** | | | |
| Distance from the City Centre | P < 0.001, $\eta^2_P$ = 0.355 | P = 0.912, $\eta^2_P$ < 0.001 | P < 0.001, $\eta^2_P$ = 0.362 |
| Human Influence Index | P < 0.001, $\eta^2_P$ = 0.276 | P = 0.767, $\eta^2_P$ < 0.001 | P < 0.001, $\eta^2_P$ = 0.034 |
| Mean Impervious Surface Cover | P < 0.001, $\eta^2_P$ = 0.313 | P = 0.901, $\eta^2_P$ < 0.001 | P < 0.001, $\eta^2_P$ = 0.136 |
| ***Desulfocapsa*** | | | |
| Distance from the City Centre | P < 0.001, $\eta^2_P$ = 0.175 | P = 0.999, $\eta^2_P$ < 0.001 | P = 0.005, $\eta^2_P$ = 0.011 |
| Human Influence Index | P < 0.001, $\eta^2_P$ = 0.175 | P = 0.971, $\eta^2_P$ < 0.001 | P = 0.005, $\eta^2_P$ = 0.011 |
| Mean Impervious Surface Cover | P < 0.001, $\eta^2_P$ = 0.176 | P = 0.999, $\eta^2_P$ < 0.001 | P = 0.024, $\eta^2_P$ < 0.001 |
| ***Desulfosporosinus*** | | | |
| Distance from the City Centre | P < 0.001, $\eta^2_P$ = 0.430 | P = 0.704, $\eta^2_P$ < 0.001 | P = 0.034, $\eta^2_P$ = 0.015 |
| Human Influence Index | P < 0.001, $\eta^2_P$ = 0.451 | P = 0.633, $\eta^2_P$ < 0.001 | P = 0.013, $\eta^2_P$ = 0.051 |
| Mean Impervious Surface Cover | P < 0.001, $\eta^2_P$ = 0.510 | P = 0.581, $\eta^2_P$ < 0.001 | P < 0.001, $\eta^2_P$ = 0.214 |
| ***Desulfuromonas*** | | | |
| Distance from the City Centre | P < 0.001, $\eta^2_P$ = 0.367 | P = 0.892, $\eta^2_P$ < 0.001 | P < 0.001, $\eta^2_P$ = 0.035 |
| Human Influence Index | P < 0.001, $\eta^2_P$ = 0.340 | P = 0.909, $\eta^2_P$ < 0.001 | P = 0.001, $\eta^2_P$ = 0.017 |
| Mean Impervious Surface Cover | P < 0.001, $\eta^2_P$ = 0.351 | P = 0.920, $\eta^2_P$ < 0.001 | P < 0.001, $\eta^2_P$ = 0.025 |
| ***Devosia*** | | | |
| Distance from the City Centre | P < 0.001, $\eta^2_P$ = 0.851 | P = 0.903, $\eta^2_P$ < 0.001 | P = 0.662, $\eta^2_P$ < 0.001 |
| Human Influence Index | P < 0.001, $\eta^2_P$ = 0.853 | P = 0.215, $\eta^2_P$ < 0.001 | P = 0.656, $\eta^2_P$ < 0.001 |
| Mean Impervious Surface Cover | P < 0.001, $\eta^2_P$ = 0.851 | P = 0.483, $\eta^2_P$ < 0.001 | P = 0.818, $\eta^2_P$ < 0.001 |
| ***Nitrosomonas*** | | | |
| Distance from the City Centre | P < 0.001, $\eta^2_P$ = 0.626 | P = 0.531, $\eta^2_P$ < 0.001 | P < 0.001, $\eta^2_P$ = 0.222 |
| Human Influence Index | P < 0.001, $\eta^2_P$ = 0.186 | P = 0.821, $\eta^2_P$ < 0.001 | P = 0.003, $\eta^2_P$ = 0.013 |
| Mean Impervious Surface Cover | P = 0.004, $\eta^2_P$ = 0.117 | P = 0.860, $\eta^2_P$ < 0.001 | P = 0.001, $\eta^2_P$ = 0.017 |
| ***Nitrospira*** | | | |
| Distance from the City Centre | P < 0.001, $\eta^2_P$ = 0.844 | P = 0.789, $\eta^2_P$ < 0.001 | P < 0.001, $\eta^2_P$ = 0.252 |
| Human Influence Index | P < 0.001, $\eta^2_P$ = 0.794 | P = 0.901, $\eta^2_P$ < 0.001 | P < 0.001, $\eta^2_P$ = 0.178 |
| Mean Impervious Surface Cover | P < 0.001, $\eta^2_P$ = 0.751 | P = 0.868, $\eta^2_P$ < 0.001 | P < 0.001, $\eta^2_P$ = 0.033 |
| ***Nitrosospira*** | | | |
| Distance from the City Centre | P < 0.001, $\eta^2_P$ = 0.786 | P = 0.728, $\eta^2_P$ < 0.001 | P < 0.001, $\eta^2_P$ = 0.154 |
| Human Influence Index | P < 0.001, $\eta^2_P$ = 0.718 | P = 0.910, $\eta^2_P$ < 0.001 | P = 0.049, $\eta^2_P$ = 0.004 |
| Mean Impervious Surface Cover | P < 0.001, $\eta^2_P$ = 0.751 | P = 0.791, $\eta^2_P$ < 0.001 | P < 0.001, $\eta^2_P$ = 0.025 |
| ***Novosphingobium*** | | | |
| Distance from the City Centre | P < 0.001, $\eta^2_P$ = 0.815 | P = 0.068, $\eta^2_P$ = 0.003 | P = 0.818, $\eta^2_P$ < 0.001 |
| Human Influence Index | P < 0.001, $\eta^2_P$ = 0.808 | P = 0.966, $\eta^2_P$ < 0.001 | P = 0.784, $\eta^2_P$ < 0.001 |
| Mean Impervious Surface Cover | P < 0.001, $\eta^2_P$ = 0.809 | P = 0.254, $\eta^2_P$ < 0.001 | P = 0.829, $\eta^2_P$ < 0.001 |
| ***Paracoccus*** | | | |
| Distance from the City Centre | P = 0.012, $\eta^2_P$ = 0.090 | P = 0.002, $\eta^2_P$ = 0.015 | P = 0.392, $\eta^2_P$ < 0.001 |
| Human Influence Index | P = 0.015, $\eta^2_P$ = 0.084 | P = 0.017, $\eta^2_P$ = 0.007 | P = 0.464, $\eta^2_P$ < 0.001 |
| Mean Impervious Surface Cover | P = 0.009, $\eta^2_P$ = 0.097 | P < 0.001, $\eta^2_P$ = 0.027 | P = 0.282, $\eta^2_P$ < 0.001 |

| Table S7 *(continued)* | | | |
|---|---|---|---|
| ***Phyllobacterium*** | | | |
| Distance from the City Centre | $P < 0.001, \eta^2_P = 0.365$ | $P = 0.225, \eta^2_P < 0.001$ | $P = 0.905, \eta^2_P < 0.001$ |
| Human Influence Index | $P < 0.001, \eta^2_P = 0.382$ | $P = 0.016, \eta^2_P = 0.007$ | $P = 0.695, \eta^2_P < 0.001$ |
| Mean Impervious Surface Cover | $P < 0.001, \eta^2_P = 0.380$ | $P = 0.024, \eta^2_P = 0.006$ | $P = 0.566, \eta^2_P < 0.001$ |
| ***Sphingobium*** | | | |
| Distance from the City Centre | $P < 0.001, \eta^2_P = 0.525$ | $P = 0.152, \eta^2_P = 0.001$ | $P = 0.995, \eta^2_P < 0.001$ |
| Human Influence Index | $P < 0.001, \eta^2_P = 0.527$ | $P = 0.113, \eta^2_P = 0.002$ | $P = 0.933, \eta^2_P < 0.001$ |
| Mean Impervious Surface Cover | $P < 0.001, \eta^2_P = 0.519$ | $P = 0.932, \eta^2_P < 0.001$ | $P = 0.933, \eta^2_P < 0.001$ |
| ***Sphingopyxis*** | | | |
| Distance from the City Centre | $P < 0.001, \eta^2_P = 0.556$ | $P = 0.187, \eta^2_P = 0.001$ | $P = 0.789, \eta^2_P < 0.001$ |
| Human Influence Index | $P < 0.001, \eta^2_P = 0.552$ | $P = 0.632, \eta^2_P < 0.001$ | $P = 0.708, \eta^2_P < 0.001$ |
| Mean Impervious Surface Cover | $P < 0.001, \eta^2_P = 0.565$ | $P = 0.056, \eta^2_P = 0.003$ | $P = 0.833, \eta^2_P < 0.001$ |
| ***Streptomyces*** | | | |
| Distance from the City Centre | $P < 0.001, \eta^2_P = 0.537$ | $P = 0.105, \eta^2_P = 0.002$ | $P = 0.860, \eta^2_P < 0.001$ |
| Human Influence Index | $P < 0.001, \eta^2_P = 0.528$ | $P = 0.694, \eta^2_P < 0.001$ | $P = 0.938, \eta^2_P < 0.001$ |
| Mean Impervious Surface Cover | $P < 0.001, \eta^2_P = 0.528$ | $P = 0.822, \eta^2_P < 0.001$ | $P = 0.997, \eta^2_P < 0.001$ |

### Supplemental Figures

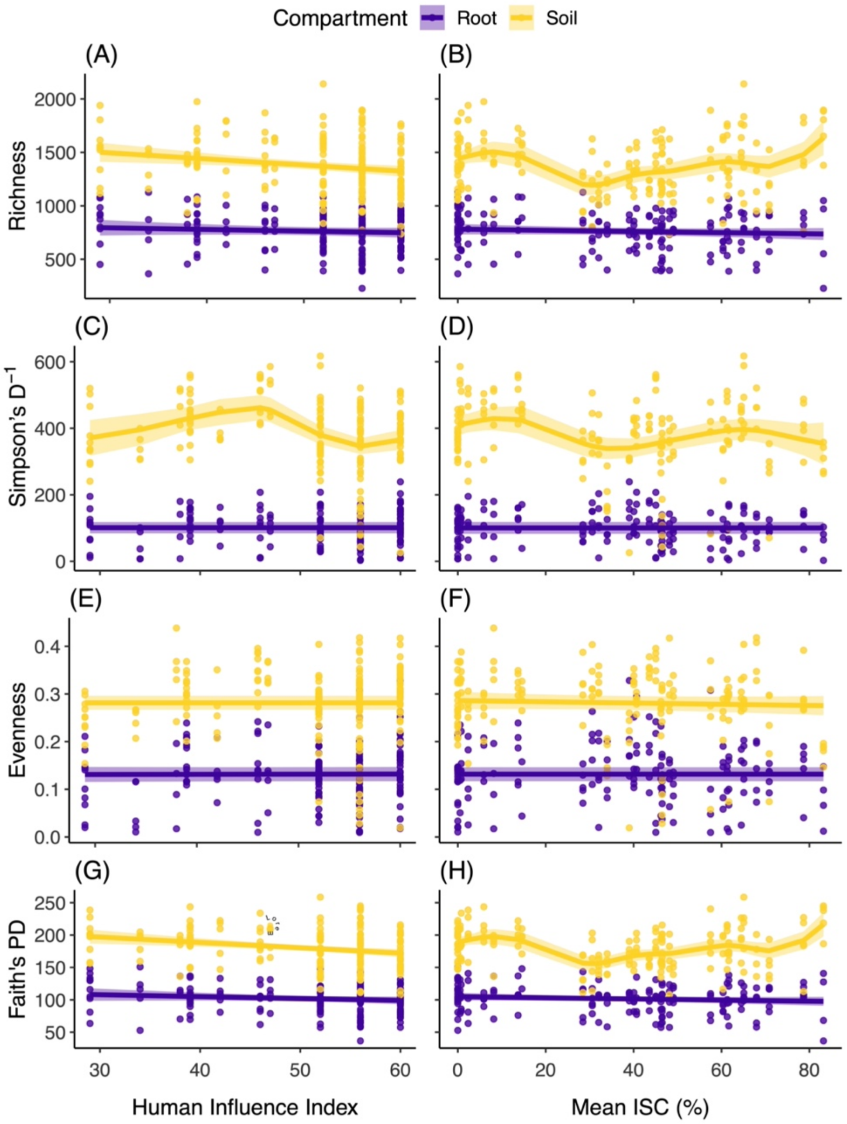


**Figure S1:** Relationship between measures of alpha diversity and Human Influence Index (A, C, E, G) and mean impervious surface cover (Mean ISC; B, D, F, H). Alpha diversity was quantified as richness (A-B), Simpson's $D^{-1}$ (C-D), evenness (E-F), and Faith's phylogenetic diversity (G-H). Points represent individual samples and lines are smoothed curves (± 95% confidence interval) from generalized additive mixed models, with separate curves shown for the root and soil microbiome compartments (purple and yellow, respectively). Deviance explained by the Human Influence Index models: richness (68.1%), Simpson's $D^{-1}$ (83.8%), evenness (66.1%), and Faith's phylogenetic diversity (73.5%). Deviance explained by the mean ISC models: richness (69.0%), Simpson's $D^{-1}$ (83.6%), evenness (66.1%), and Faith's phylogenetic diversity (74.4%). Detailed statistics are provided in Table S2.

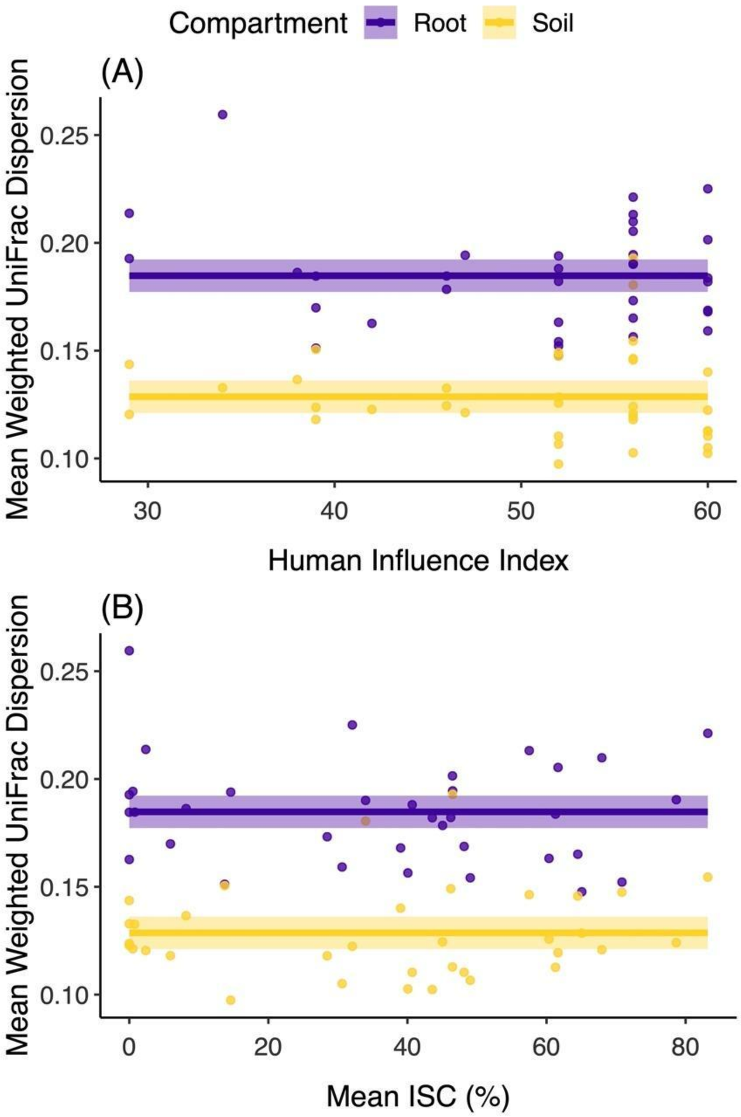


**Figure S2:** Relationship between mean weighted UniFrac beta dispersion and (A) Human Influence Index and (B) mean impervious surface cover (mean ISC). Points represent population means and lines are smoothed curves (± 95% confidence interval) from a generalized additive model, with separate curves shown for the root and soil microbiome compartments (purple and yellow, respectively). Deviance explained by the models: Human Influence Index = 60.5% and mean ISC = 61.0%. Detailed statistics are provided in Table S4.

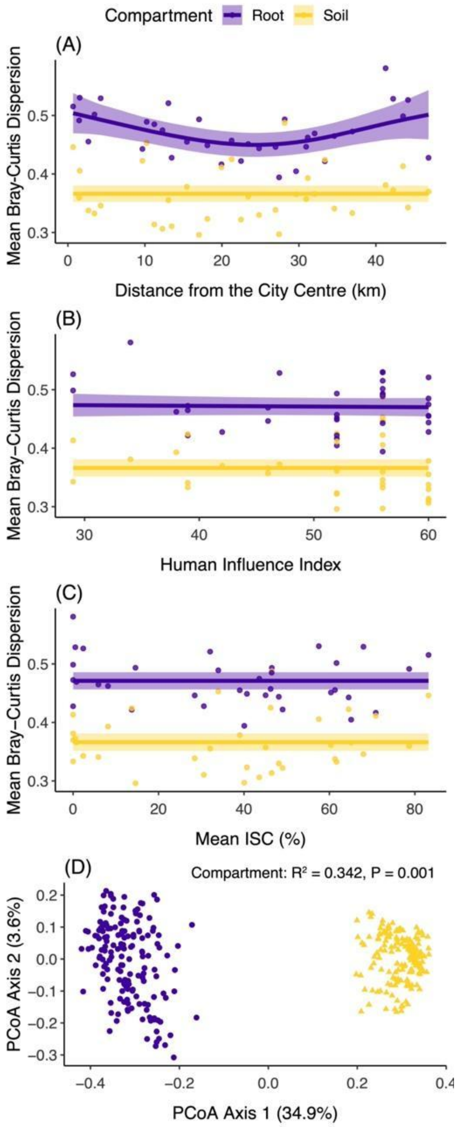


**Figure S3:** Relationship between mean Bray-Curtis beta dispersion and (A) distance from the city centre, (B) Human Influence Index, and (C) mean impervious surface cover (mean ISC). Points represent population means and lines are smoothed curves (± 95% confidence interval) from a generalized additive model, with separate curves shown for the root and soil microbiome compartments (purple and yellow, respectively). Deviance explained by the models: distance from the city centre = 64.7%, Human Influence Index = 58.9%, and mean ISC = 58.7%. Detailed statistics are provided in Table S3. We also show the principal coordinates analysis of the Bray-Curtis dissimilarity among bacterial communities (D), which shows distinct groupings by microbiome compartment (purple circles = root, yellow triangles = soil). Inset text presents results from a PERMANOVA comparing composition by microbiome compartment.

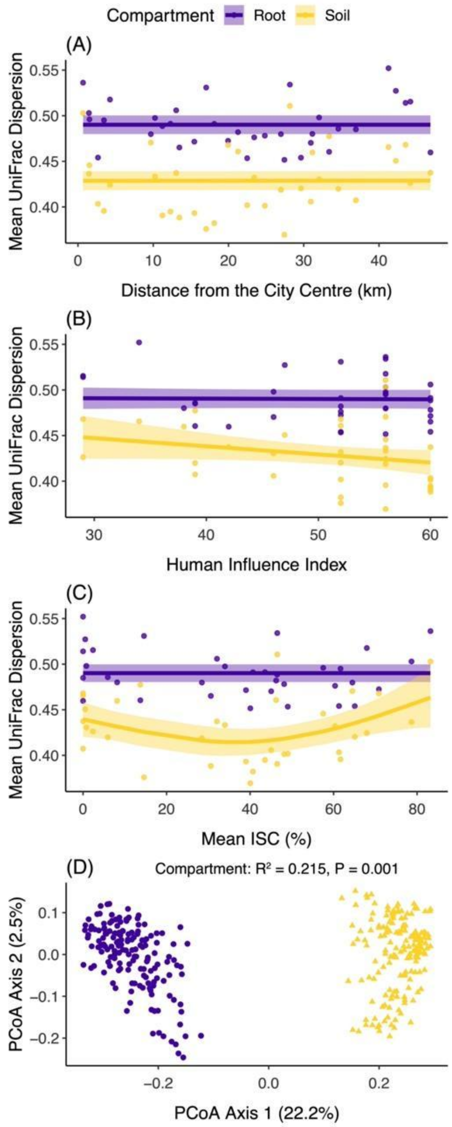


**Figure S4:** Relationship between mean UniFrac beta dispersion and (A) distance from the city centre, (B) Human Influence Index, and (C) mean impervious surface cover (mean ISC). Points represent population means and lines are smoothed curves (± 95% confidence interval) from a generalized additive model, with separate curves shown for the root and soil microbiome compartments (purple and yellow, respectively). Deviance explained by the models: distance from the city centre = 50.2%, Human Influence Index = 53.0%, and mean ISC = 57.0%. Detailed statistics are provided in Table S3. We also show the principal coordinates analysis of the UniFrac dissimilarity among bacterial communities (D), which shows distinct groupings by microbiome compartments (purple circles = root, yellow triangles = soil). Inset text presents results from a PERMANOVA comparing composition by microbiome compartment.

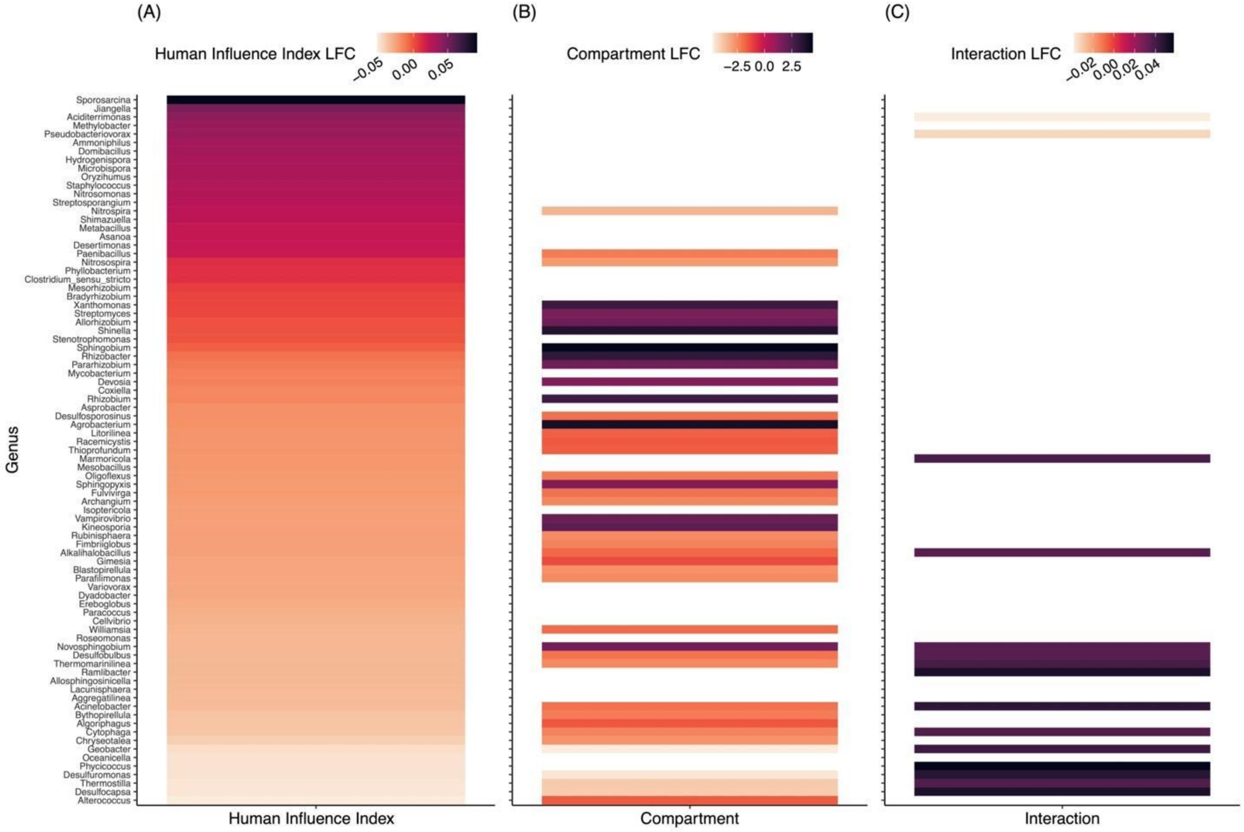


**Figure S5:** Heatmaps of differentially-abundant bacterial taxa by Human Influence Index (A), compartment (root or soil; B) and the interaction (C) following the ANCOM-BC2 procedure. For simplicity, we show taxa with $P < 0.100$ after false discovery rate correction, with taxa ordered in descending log-fold change (LFC) in relation to the Human Influence Index. We set soil as the reference compartment, and positive LFCs indicate increased abundance with increasing human influence (i.e., greater anthropogenic impacts, A), increased abundance in the root compartment (B), and increased abundance in the root compartment with increasing human influence. Full results of the differential abundance analysis are provided in a spreadsheet in the electronic supplementary material. Note: the LFC scale is different for each panel.

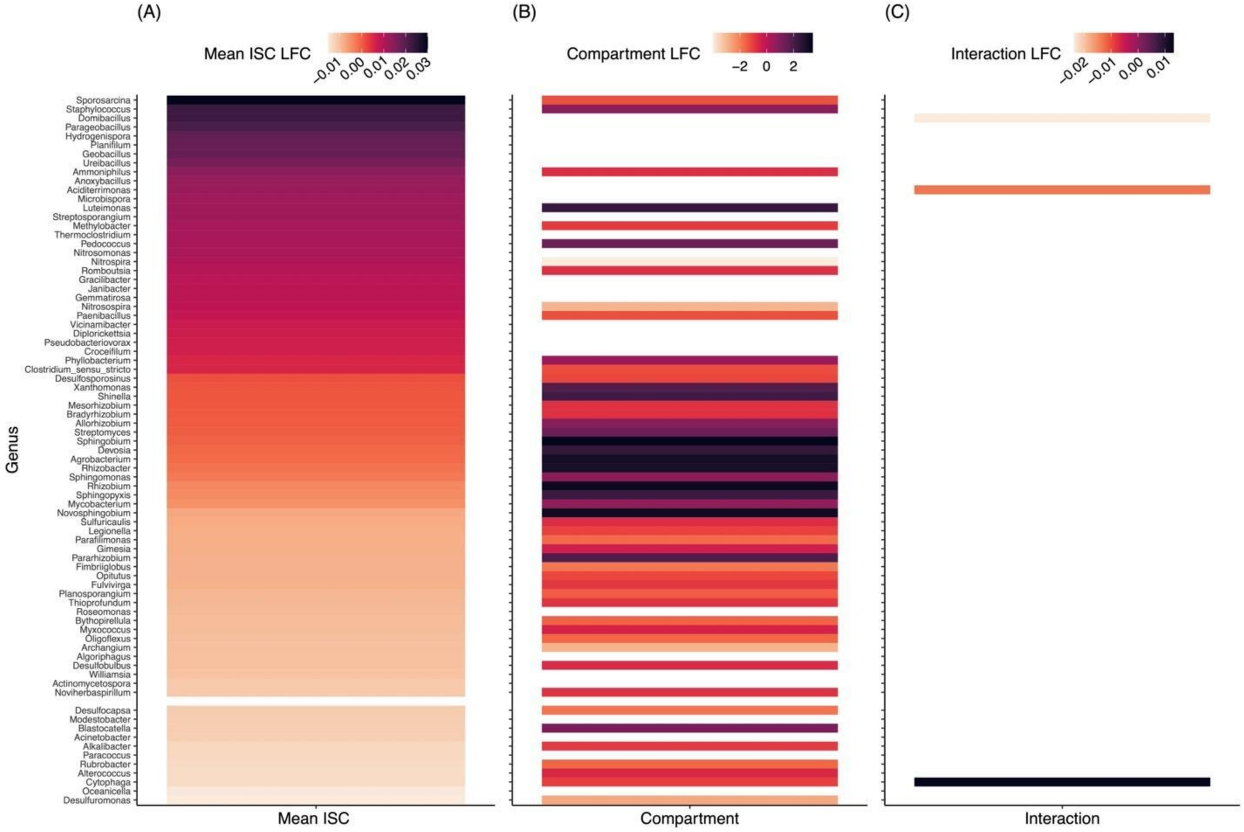


**Figure S6:** Heatmaps of differentially-abundant bacterial taxa by mean impervious surface cover (mean ISC; A), compartment (root or soil; B) and the interaction (C) following the ANCOM-BC2 procedure. For simplicity, we show taxa with $P < 0.100$ after false discovery rate correction, with taxa ordered in descending log-fold change (LFC) in relation to mean ISC. We set soil as the reference compartment, and positive LFCs indicate increased abundance with increasing ISC (A), increased abundance in the root compartment (B), and increased abundance in the root compartment with increasing ISC. Full results of the differential abundance analysis are provided in a spreadsheet in the electronic supplementary material. Note: the LFC scale is different for each panel.

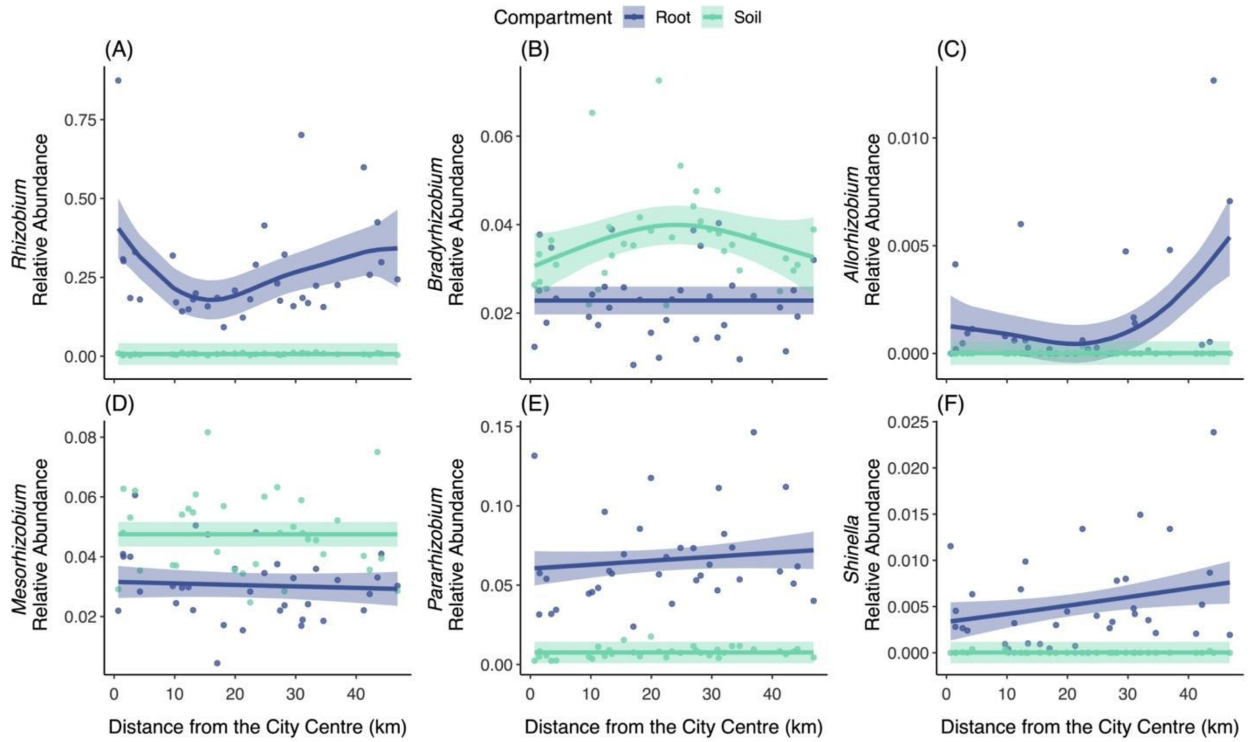


**Figure S7:** Relationships between the relative abundance of focal mutualistic bacteria taxa and distance from the city centre. Points represent population means and lines are smoothed curves (± 95% confidence interval) from generalized additive models, with separate curves shown for the root and soil microbiome compartments (blue and green, respectively). Deviance explained by the models: *Rhizobium* (66.8%), *Bradyrhizobium* (39.7%), *Allorhizobium* (36.9%), *Mesorhizobium* (33.2%), *Pararhizobium* (68.3%), and *Shinella* (40.8%). Detailed statistics are provided in Table S6. Note: the scale of the y-axis varies by focal bacterium.

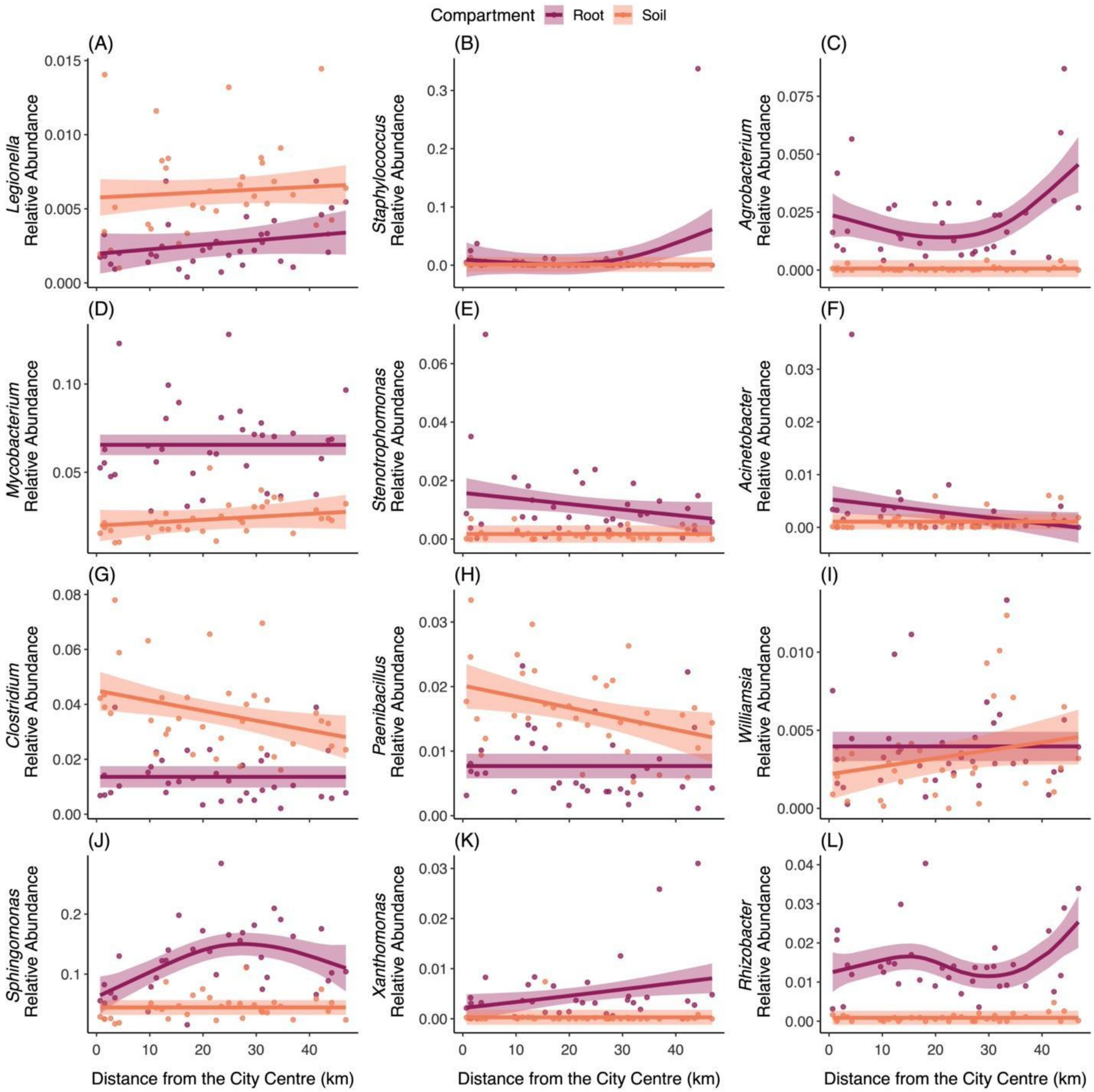


**Figure S8:** Relationships between the relative abundance of focal pathogenic bacteria taxa and distance from the city centre. Points represent population means and lines are smoothed curves (± 95% confidence interval) from generalized additive models, with separate curves shown for the root and soil microbiome compartments (red and orange, respectively). Deviance explained by the models: *Legionella* (49.4%), *Staphylococcus* (15.4%), *Agrobacterium* (56.4%), *Mycobacterium* (60.1%), *Stenotrophomonas* (27.6%), *Acinetobacter* (11.4%), *Clostridium* (53.4%), *Paenibacillus* (41.3%), *Williamsia* (6.2%), *Sphingomonas* (60.4%), *Xanthomonas* (27.1%), and *Rhizobacter* (67.6%). Detailed statistics are provided in Table S7. Note: the scale of the y-axis varies by focal bacterium.

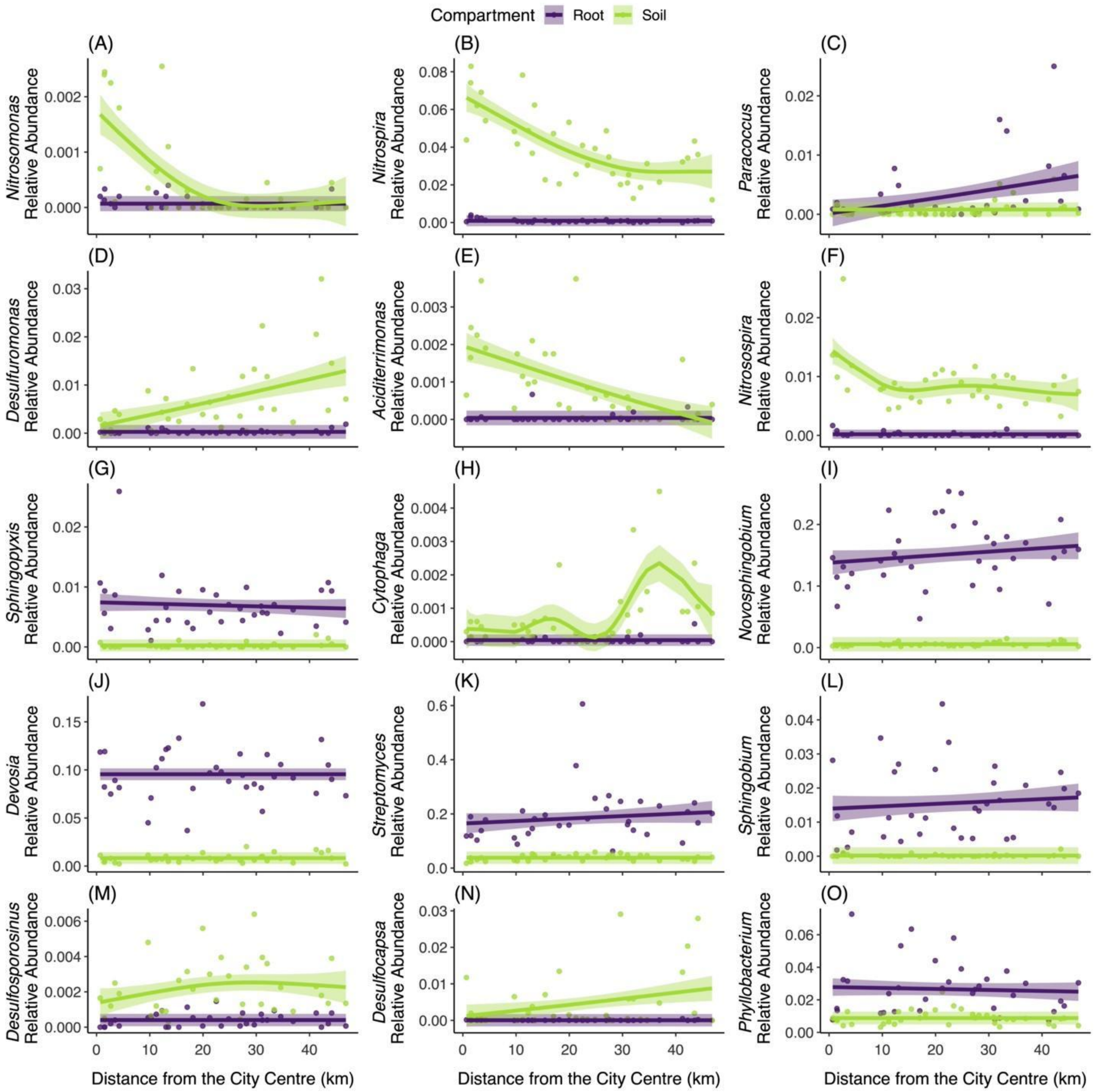


**Figure S9:** Relationships between the relative abundance of focal ecosystem function bacteria taxa and distance from the city centre. Points represent population means and lines are smoothed curves (± 95% confidence interval) from generalized additive models, with separate curves shown for the root and soil microbiome compartments (purple and green, respectively). Deviance explained by the models: *Nitrosomonas* (55.5%), *Nitrospira* (86.8%), *Paracoccus* (20.4%), *Desulfuromonas* (47.7%), *Aciditerrimonas* (54.6%), *Nitrosospira* (80.8%), *Sphingopyxis* (56.1%), *Cytophaga* (59.9%), *Novosphingobium* (81.7%), *Devosia* (85.1%), *Streptomyces* (54.5%), *Sphingobium* (53.0%), *Desulfosporosinus* (46.4%), *Desulfocapsa* (25.1%), and *Phyllobacterium* (37.0%). Detailed statistics are provided in Table S8. Note: the scale of the y-axis varies by focal bacterium.

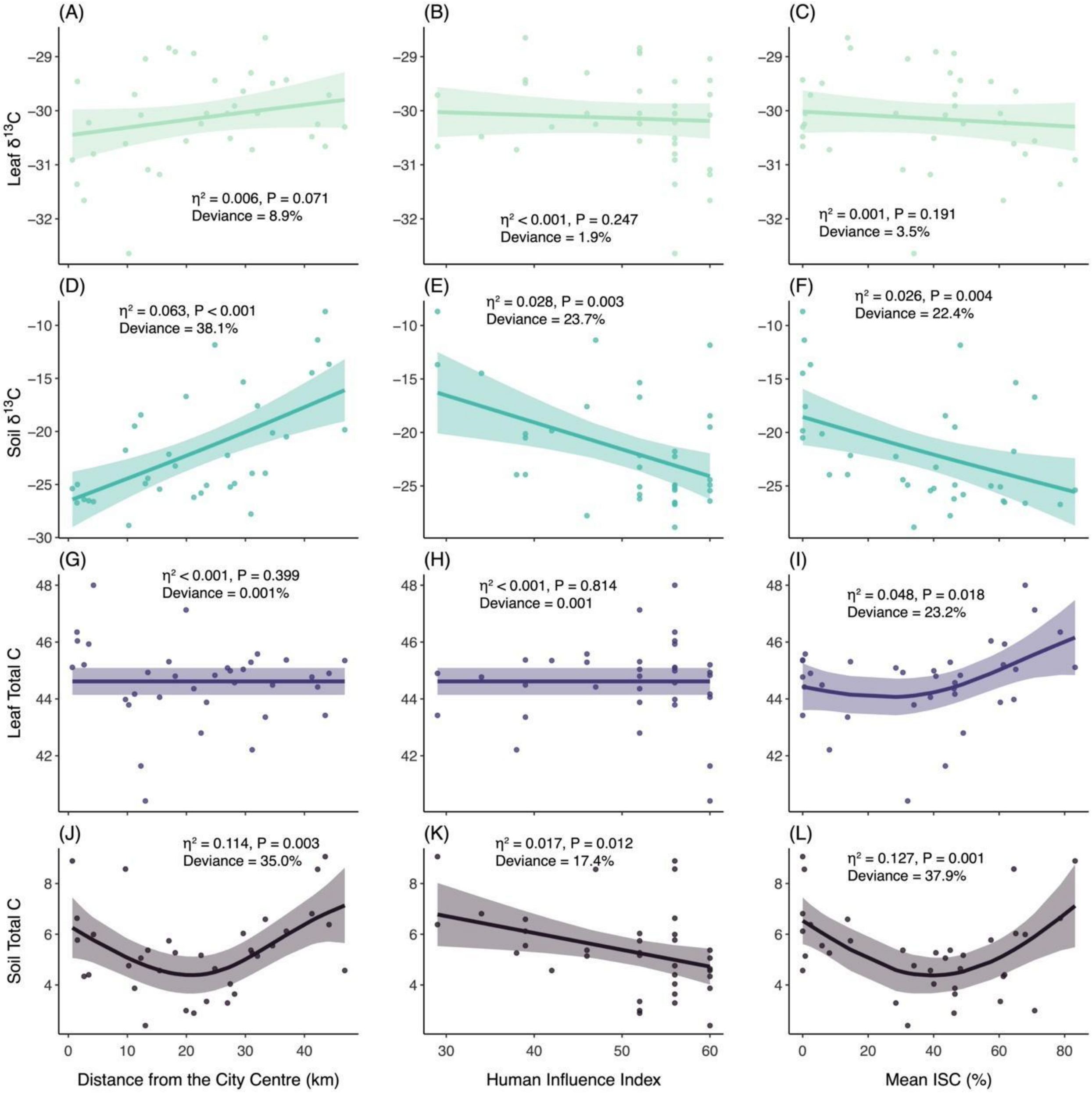


**Figure S10:** Plots of leaf $\delta^{13}C$ (A-C), soil $\delta^{13}C$ (D-F), leaf total C (G-I), and soil total C (J-L) against distance from the city centre (A, D, G, J), Human Influence Index (B, E, H, K), and mean impervious surface cover (mean ISC; C, F, I, L). Points represent population means and lines are smoothed curves (± 95% confidence interval) from generalized additive models. Colours indicate different metrics (leaf $\delta^{13}C$ = green, soil $\delta^{13}C$ = blue, leaf total C = purple, and soil total C = black). Inset text provides effect size (eta squared, $\eta^2$) and P-value for the smoothed curve, with the deviance explained by the model also provided.

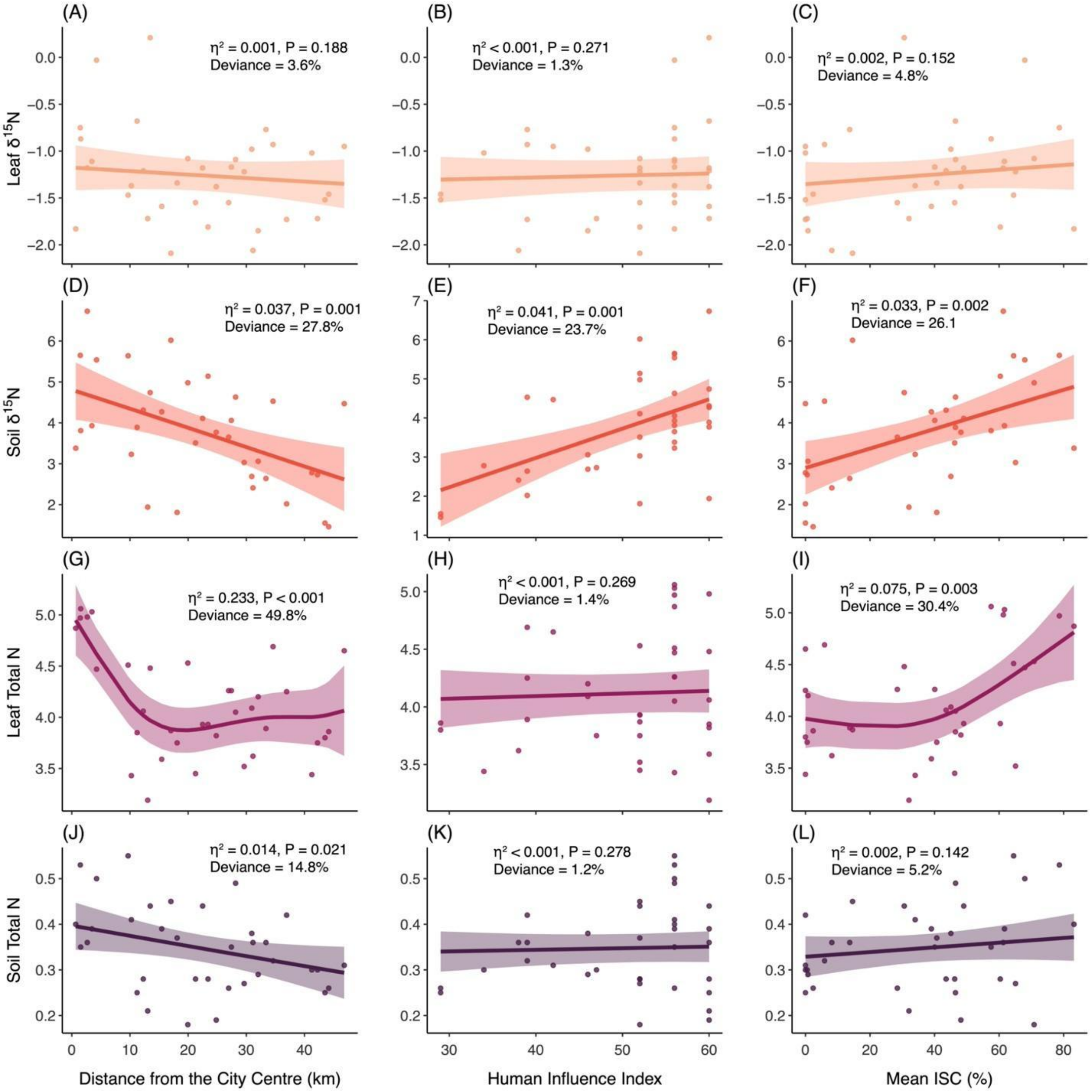


**Figure S11:** Plots of leaf $\delta^{15}N$ (A-C), soil $\delta^{15}N$ (D-F), leaf total N (G-I), and soil total N (J-L) against distance from the city centre (A, D, G, J), Human Influence Index (B, E, H, K), and mean impervious surface cover (mean ISC; C, F, I, L). Points represent population means and lines are smoothed curves (± 95% confidence interval) from generalized additive models. Colours indicate different metrics (leaf $\delta^{15}N$ = light orange, soil $\delta^{15}N$ = dark orange, leaf total N = dark red, and soil total N = black). Inset text provides effect size (eta squared, $\eta^2$) and P-value for the smoothed curve, with the deviance explained by the model also provided.
